# PU.1 enforces quiescence and limits hematopoietic stem cell expansion during inflammatory stress

**DOI:** 10.1101/2020.05.18.102830

**Authors:** James S. Chavez, Jennifer L. Rabe, Dirk Loeffler, Kelly C. Higa, Giovanny Hernandez, Taylor S. Mills, Nouraiz Ahmed, Rachel L. Gessner, Zhonghe Ke, Beau M. Idler, Hyun Min Kim, Jason R. Myers, Brett M. Stevens, Craig T. Jordan, Hideaki Nakajima, John Ashton, Robert S. Welner, Timm Schroeder, James DeGregori, Eric M. Pietras

**Author notes:** Correspondence should be addressed to E.M.P.

## Abstract

Loss of hematopoietic stem cell (HSC) quiescence and resulting clonal expansion are common initiating events in the development of hematological malignancy. Likewise, chronic inflammation related to aging, disease and/or tissue damage is associated with leukemia progression, though its role in oncogenesis is not clearly defined. Here, we show that PU.1-dependent repression of protein synthesis and cell cycle genes in HSC enforces homeostatic protein synthesis levels and HSC quiescence in response to IL-1 stimulation. These genes are constitutively de-repressed in PU.1-deficient HSC, leading to activation of protein synthesis, loss of quiescence and aberrant expansion of HSC. Taken together, our data identify a mechanism whereby HSC regulate their cell cycle activity and pool size in response to chronic inflammatory stress.

## Introduction

Hematopoietic stem cell (HSC) quiescence promotes lifelong blood regeneration and is controlled by a complex regulatory network including cell-intrinsic transcription factors and epigenetic modifiers (Pietras et al., 2011), organelle homeostasis mechanisms (Hinge et al., 2020; Liang et al., 2020), and signals generated from the bone marrow (BM) niche (Morrison and Scadden, 2014). HSC can be briefly forced out of quiescence to facilitate blood system regeneration by numerous stressors such as, infection (Prendergast and Essers, 2014), chronic stress (Heidt et al., 2014) and myeloablative injury (Harrison and Lerner, 1991) demonstrating that HSC are responsive to disruptions in BM homeostasis (King and Goodell, 2011). HSC limit cell cycle activation in response to physiological stressors and resultant inflammatory signaling, which can impair HSC function (Pietras, 2017). While transient proliferative responses to stress are crucial adaptations that promote organismal survival, long-term loss of HSC quiescence can lead to bone marrow failure, overproduction of potentially toxic cell types and/or hematological malignancy (Pietras, 2017). At least a small HSC fraction can remain quiescent despite exposure to stressors like myeloablation, functioning in a ‘reserve’ capacity to maintain long-term blood system integrity (Wilson et al., 2008; Zhao et al., 2019).

HSC can directly respond to pathogens and physiological ‘danger’ signals via direct sensing but may also respond indirectly via paracrine pro-inflammatory cytokine signaling (Takizawa et al., 2017). Pro-inflammatory cytokine induction, which is common amongst nearly all physiological stressors, can drive HSC proliferation or differentiation (Mirantes et al., 2014). Notably, HSC proliferate in response to IFNs (Ehninger et al., 2014; Essers et al., 2009; Matatall et al., 2016), LPS (Takizawa et al., 2017), G-CSF (Schuettpelz et al., 2014), TNF (Yamashita and Passegue, 2019) and interleukin (IL)-1 (Hemmati et al., 2019; Weisser et al., 2016). IL-1 consists of two cytokines (IL-1 α and β) with different expression patterns that share a common receptor complex and elicit similar responses (Dinarello, 2018). IL-1 is produced in response to a wide variety of physiological stress conditions including aging, chronic inflammatory disease, myeloablative treatments such as radiation and/or chemotherapy, obesity and cellular senescence (Dinarello, 2018; Laberge et al., 2015). It may also contribute to hematological malignancy and is highly expressed in cells from patients with myelodysplastic syndrome (MDS), myeloproliferative neoplasia (MPN), and acute myelogenous leukemia (AML) (Agerstam et al., 2016; Barreyro et al., 2018; Carey et al., 2017; Ezaki et al., 1995; Zhang et al., 2016). Previous work from our group demonstrates that acute IL-1 exposure drives myeloid cell overproduction *in vivo*, via precocious activation of the master myeloid transcription factor PU.1 (Pietras et al., 2016) in HSC. Interestingly, this effect was transient and HSC re-entered quiescence following chronic exposure to IL-1 suggesting the existence of a ‘braking’ mechanism that limits HSC cell cycle entry (Pietras et al., 2016; Rabe et al., 2020).

We recently identified a broad downregulation of cell proliferation genes in quiescent HSC from a mouse model of rheumatoid arthritis depending in part on IL-1 signaling (Hernandez et al., 2020). To further explore the relevance of this gene program to HSC function during chronic inflammation, we used a mouse model of chronic IL-1 stimulation (Pietras et al., 2016). We show that IL-1 is sufficient to induce repression of a broad set of cell cycle and protein synthesis genes, in turn limiting HSC cell cycle activation. We find that repression of these genes in HSC is associated with high PU.1 levels and ChIP-seq analysis shows that PU.1 directly binds these targets. Strikingly, genetic reduction of PU.1 levels increases expression of cell cycle and protein synthesis genes, licensing HSC to increase protein synthesis activity and enter the cell cycle in response to IL-1. We find these effects are coupled with excess myeloid lineage expansion reminiscent of early hematological malignancy (Sica et al., 2019). Together, these data support a model in which PU.1 enforces HSC quiescence in response to inflammatory stress by limiting protein synthesis and cell cycle activity, thereby regulating the size of the HSC pool.

## Results

### Chronic IL-1 induces repression of cell cycle and protein synthesis genes

HSC maintain a largely quiescent state under inflammatory stress (Hernandez et al., 2020; Pietras, 2017; Pietras et al., 2014; Rabe et al., 2020). Interestingly, our molecular profiling of HSC in the collagen-induced arthritis (CIA) mouse model of rheumatoid arthritis (RA) showed HSC quiescence was associated with repression of genes associated with cell proliferation and protein synthesis pathways, such as *Myc* (Hernandez et al., 2020; Rabe et al., 2020). To address the underlying mechanism(s), we analyzed gene expression by RNA-seq in the HSC-enriched SLAM fraction (SLAM cells; LSK/Flk2^-^CD48^-^CD150^+^) isolated from the BM of mice treated ± interleukin-1β (IL-1) for 20 days to model chronic inflammatory stress (Pietras et al., 2016; Rabe et al., 2020) (**Fig. 1A**). We found over 1400 differentially expressed genes (DEGs; padj <0.05) (**Fig. 1B, Table S1**). Gene ontology (GO) analysis identified *cell activation, immune response, leukocyte adhesion* and *defense response* gene programs among upregulated DEGs (**Fig. S1A, Table S2**). Using a custom Fluidigm qRT-PCR array, we validated increased expression of key genes in these pathways including *Itgam* (Mac-1), *Cdk6, Flt3, Il1r1, Slamf1* (CD150) and *Pdgfrb* (**Fig. S1B**). Strikingly, downregulated DEGs were enriched for *ribosome biogenesis, rRNA processing, tRNA processing*, and *translation* GO categories (**Fig. 1C, Table S2**), while Ingenuity Pathway Analysis (IPA) and GSEA likewise identified multiple cell proliferation and mRNA translation pathways inhibited by IL-1 (**Fig. 1D-E, Fig. S1C, Table S2**). We also validated these results by Fluidigm qRT-PCR, confirming that *Myc* and several other genes regulating rRNA synthesis and HSC cell cycle entry were downregulated, while the cell cycle inhibitor *Cdkn1b* was upregulated (**Fig. S1D**). These data were strikingly reminiscent of *Myc* pathway downregulation we observed in HSC from mice with CIA (Hernandez et al., 2020), suggesting they are not simply a byproduct of *in vivo* treatment with IL-1.

**Figure 1.**
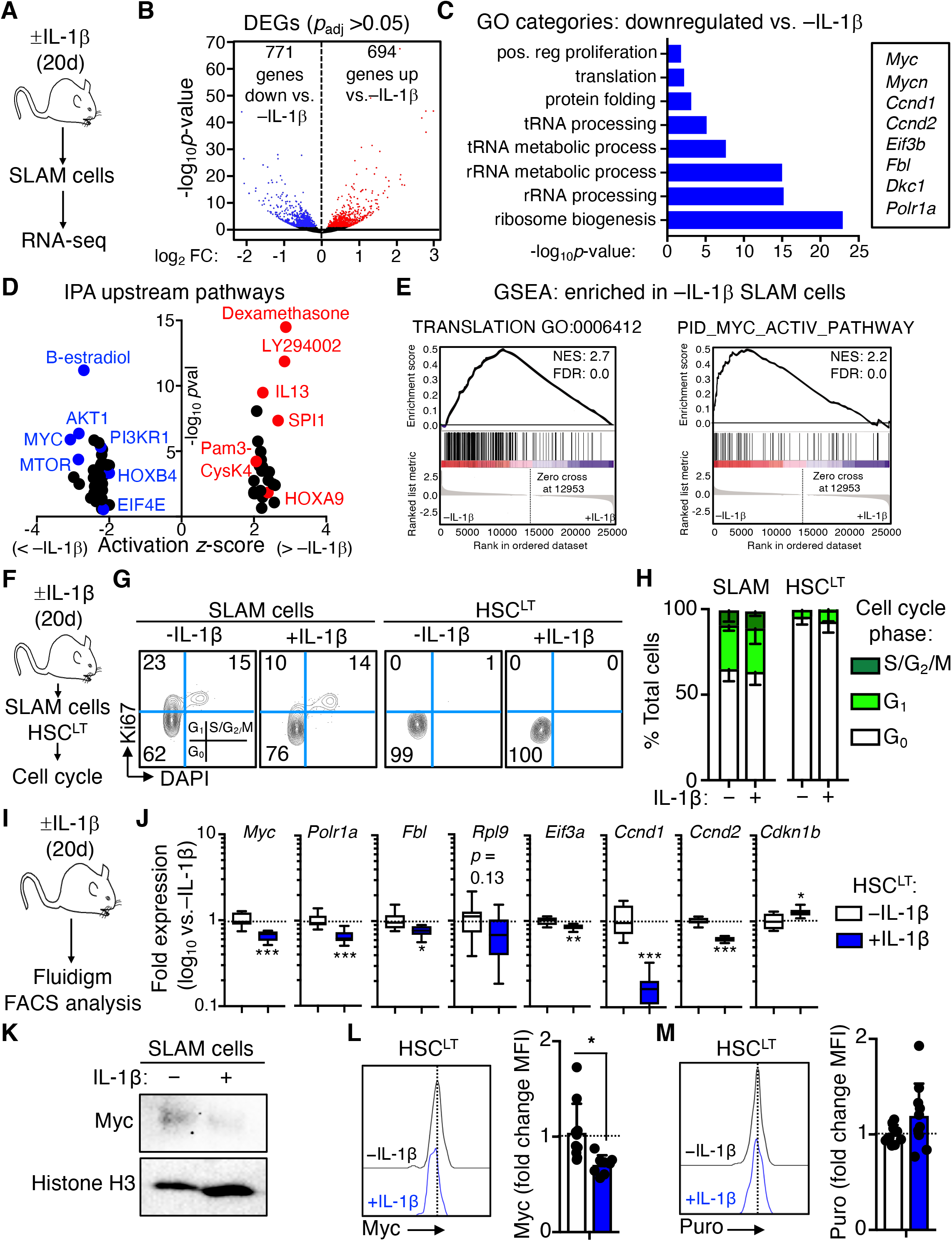
Chronic IL-1 induces repression of cell cycle and protein synthesis genes. **A**) Experimental design for RNA-seq studies. (n = 4-7 pools of SLAM cells) from mice treated for 20d ± IL-1. **B**) Volcano plot of differentially expressed genes (padj ≤ 0.05) in IL-1-exposed SLAM cells from A) showing log_2_ fold change (FC) versus −log_10_ p-value significance. See also Table S1. **C**) GO category enrichment of downregulated DEGs in IL-1-exposed SLAM cells from A), expressed as −log_10_ p-value. See also Table S2. **D**) Ingenuity Pathway Analysis (IPA) showing enriched upstream regulators of DEGs in IL-1-exposed SLAM cells from A). See also Table S3. **E**) GSEA analysis of significantly downregulated DEGs. GSEA plots show negative enrichment of translation and Myc pathway genes in IL-1-exposed SLAM cells from A). See also Table S4. **F**) Experimental design for cell cycle analyses of SLAM cells and HSC^LT^ from mice treated for 20d ±IL-1. **G**) Representative flow cytometry plots showing cell cycle distribution in SLAM cells and HSC^LT^ from F). **H**) Quantification of cell cycle phase distribution in SLAM cells and HSC^LT^ from F) (n = 5/group). Data are from one experiment. **I**) Experimental design for Fluidigm qRT-PCR analyses and intracellular FACS staining HSC^LT^ of SLAM cells and HSC^LT^ from mice treated for 20d ± IL-1. **J**) Quantification by Fluidigm qRT-PCR array of cell cycle and protein synthesis gene expression in HSC^LT^ from I) (n = 8/group). Data are expressed as log_10_ fold expression versus-IL-1β. Box represents upper and lower quartiles with line representing median value. Whiskers represent minimum and maximum values. Data are representative of two independent experiments. **K**) Western Blot analysis of Myc levels in SLAM cells from mice treated ± IL-1β for 20d. Histone H3 loading control levels from the same blot is shown. Data are from one experiment. **L**) Intracellular flow cytometry analysis of Myc protein levels in HSC^LT^ (n = 10 -IL-1β, 8 +IL-1β). Data are expressed as fold change of mean fluorescence intensity (MFI) versus -IL-1β. Individual values are shown with bars representing mean values. Data are compiled from three independent experiments. **M**) Intracellular flow cytometry analysis of puromycin (Puro) incorporation in HSC^LT^ (n = 9 -IL-1β, 10 +IL-1β). Data are expressed as fold change of mean fluorescence intensity (MFI) versus -IL-1β. Individual values are shown with bars representing mean values. Data are compiled from three independent experiments. * p< 0.05, ** p< 0.01, ***p< 0.001 by Mann-Whitney *u*-test. Error bars represent S.D. See also Figure S1.

We previously showed that long-term HSC (HSC^LT^)-enriched SLAM cells (defined as LSK/Flk2^-^CD48^-^CD150^+^CD34^-^EPCR^+^) overlap phenotypically with deeply quiescent CD49b^lo^ ‘reserve’ HSC (Zhao et al., 2019) and remain almost exclusively in a quiescent (G_0_) cell cycle state despite chronic IL-1 exposure (Rabe et al., 2020). We confirmed that chronic IL-1 did not increase HSC^LT^ cell cycle activity in an independent cohort of mice (**Fig. 1F-H, Fig. S1E**). Moreover, while acute (1d) IL-1 exposure activated SLAM cell proliferation, HSC^LT^ remained largely quiescent, suggesting this mechanism may be initiated rapidly upon IL-1 stimulation (**Fig. S1F-H**). Consistent with our RNA-seq data in SLAM cells, cell cycle and protein synthesis genes were repressed in chronic IL-1-exposed HSC^LT^ (**Fig. 1I-J**). We also confirmed reduced Myc expression in HSC by western blot (**Fig. 1K**) and intracellular flow cytometry staining (**Fig. 1L, Fig. S1I**). Myc levels in phenotypic MPP4 (LSK/Flk2^+^) were not significantly impacted by chronic IL-1, suggesting not all progenitor cells may repress Myc in response to IL-1 (**Fig. S1I**). Since chronic IL-1 exposure reduced expression of protein synthesis genes, we also measured the impact of chronic IL-1 exposure on protein synthesis rates in HSC^LT^ by *in vivo* puromycin (puro) incorporation assays (**Fig. 1M**). Consistent with prior work (Signer et al., 2014), puro incorporation rates in HSC^LT^, SLAM cells and MPP4 were all similar (**Fig. S1J**), and chronic IL-1 exposure did not impact the HSC^LT^ protein synthesis rate (**Fig. 1M, Fig. S1J**). Taken together these data suggest the IL-1-induced gene repression program could serve as a rheostatic mechanism to maintain homeostatic protein synthesis and cell cycle activity in HSC^LT^ during chronic inflammatory stress.

### Direct IL-1 stimulation restricts HSC^LT^ cell cycle entry

To assess the direct impact of IL-1 signaling on HSC^LT^, we analyzed the HSC^LT^ cell cycle activity and gene expression using *in vitro* culture. Notably, cell cycle entry was delayed in HSC^LT^ cultured for 24h +IL-1 in a *MyD88*-dependent fashion (**Fig. 2A**), supporting a direct signaling mechanism via the MyD88 axis downstream of IL-1R (**Fig. S2A**). We next analyzed HSC^LT^ division kinetics ± IL-1 using live single-cell imaging of cultured *PU.1-EYFP* HSC^LT^ (**Fig. 2B**). Consistent with our cell cycle analyses, IL-1 significantly delayed (but did not halt) the initial cell division of cultured HSC^LT^ (**Fig. 2C-D**), reminiscent of our prior findings in cultured SLAM cells (Pietras et al., 2016). PU.1-EYFP levels were also rapidly and significantly increased in HSC^LT^ cultured +IL-1 prior to the first cell division, also consistent with previous observations for IL-1 and TNF (Etzrodt et al., 2019) (**Fig. 2E, Fig. S2B-D**). IL-1 treatment also rapidly repressed cell cycle and protein synthesis genes in HSC^LT^ after 12h in culture (**Fig. 2F-G**), with a corresponding increase in *Spi1* and *Itgam* (**Fig. S2E**). Given the broad downregulation of protein synthesis genes, we assessed whether reduced protein synthesis is sufficient to limit HSC^LT^ cell cycle entry. We thus compared the impact of IL-1 versus omacetaxine (Oma), which binds the ribosomal A-site and inhibits protein synthesis (Gandhi et al., 2014), on HSC^LT^ cell cycle entry (**Fig. 2H**). Like IL-1, Oma effectively limited HSC cell cycle entry and reduced total Ki67 protein expression (**Fig. 2I-J**). Since reduced Ki67 levels may be related to translation inhibition and not cell cycle changes, we independently read out cell cycle progression based on the frequency of cells with >2N DNA content (ie, S/G_2_/M phases), based on DAPI alone. We observed a significant reduction in IL-1 and/or Oma-treated HSC^LT^ in S phase or beyond, consistent with slowed cell cycle activity (**Fig. 2J**). Oma and IL-1 also reduced the forward scatter (FSC) profile of cultured HSC, consistent with previous observations correlating cell size and biosynthetic activity (Iritani and Eisenman, 1999; Scognamiglio et al., 2016) (**Fig. S2F**). Collectively, these data indicate that IL-1 directly and rapidly represses cell cycle and protein synthesis genes, thereby limiting HSC^LT^ cell cycle entry.

**Figure 2.**
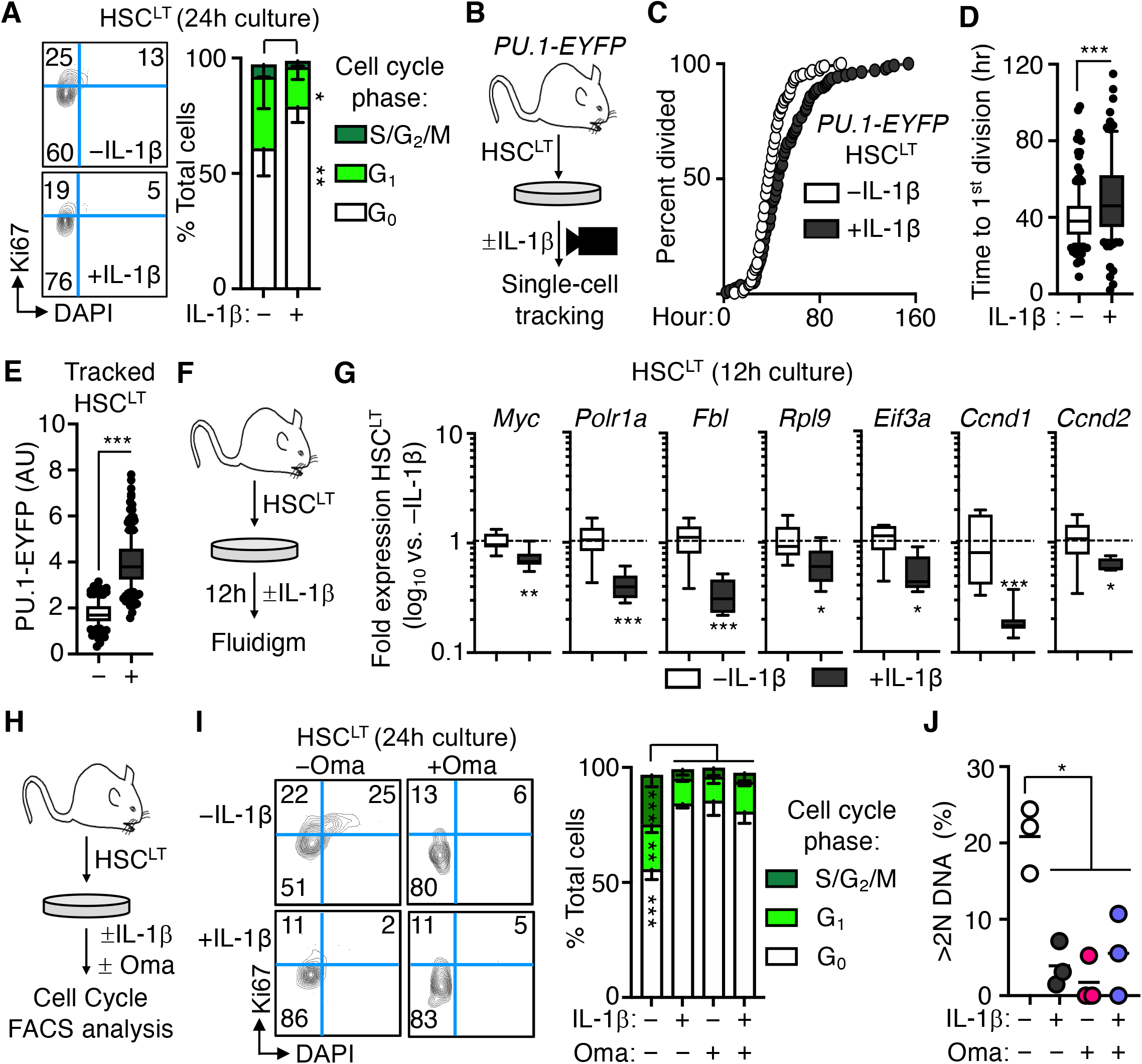
Direct IL-1 stimulation restricts HSC^LT^ cell cycle entry. **A**) Representative FACS plots (left) and quantification (right) of cell cycle distribution in HSC^LT^ cultured 24h ± IL-1 (n = 6/group). Data are compiled from three independent experiments. **B**) Experimental design for single cell tracking studies of *PU.1-EYFP* HSC^LT^ cultured ± IL-1. Time to first cell division was tracked via microscopy. **C**) Graph showing kinetics of first cell division in HSC^LT^ from B) (n = 194 -IL-1β, 139 IL-1β). Data are compiled from three independent experiments. **D**) Cumulative time to first division of HSC^LT^ from B). Data are compiled from three independent experiments. Box shows upper and lower quartiles with line showing median value; whiskers upper and lower 10^th^ percentile and individual dots represent outliers. **E**) PU.1-EYFP levels in HSC^LT^ prior to first cell division. (n = 137-IL-1β, 192 +IL-1β). Data are compiled from three independent experiments. Box shows upper and lower quartiles with line showing median value; whiskers upper and lower 10^th^ percentile and individual dots represent outliers. **F**) Experimental design for Fluidigm qRT-PCR array analysis of HSC^LT^ cultured ± IL-1β for 12h. **G**) Quantification by Fluidigm qRT-PCR array of cell cycle and protein synthesis gene expression in HSC^LT^ from F) (n = 8/group). Data are expressed as log_10_ fold expression versus-IL-1β. Box represents upper and lower quartiles with line representing median value. Whiskers represent minimum and maximum values. Data are representative of two independent experiments. **H**) Experimental design for cell cycle analysis of HSC^LT^ cultured ± IL-1 and ± 10 μM Omacetaxine (Oma) for 24h. **I**) Representative FACS plots (left) and quantification (right) of cell cycle distribution in HSC^LT^ from H) (n = 3/group). Data are from one experiment. **J**) Proportion of HSC^LT^ with >2N DAPI signal from H). Individual values are shown with means (horizontal line). * p< 0.05, ** p< 0.01, ***p< 0.001 by Mann-Whitney *u*-test or one-way ANOVA with Tukey’s test in I) and J). Error bars represent S.D. See also Figure S2.

### IL-1 induced gene repression is associated with increased PU.1 activity

To investigate how IL-1 represses cell cycle and protein synthesis genes in HSC^LT^, we reanalyzed our IPA data and noticed it had identified SPI1 (PU.1) pathway activation in IL-1-exposed SLAM cells (**Fig. 1D**), consistent with our prior published findings (Pietras et al., 2016; Rabe et al., 2020). Interestingly, PU.1 can also restrict proliferation in hematopoietic stem and progenitor cells (HSPC), likely a mechanism to promote PU.1 accumulation during myeloid differentiation (Fukuchi et al., 2008; Kueh et al., 2013). We reasoned this mechanism could also serve to restrict HSC activation by IL-1. To establish a relationship between PU.1 and IL-1-mediated repression of cell cycle and protein synthesis genes, we first compared our RNA-seq dataset with publicly available datasets in which PU.1 was overexpressed in thymocytes ((Hosokawa et al., 2018); GSE93755) and in which *Myc* and *Mycn* were conditionally deleted in HSC, leading to reduced proliferative activity ((Laurenti et al., 2008); GSE12467). GSEA analysis revealed that cell cycle and protein synthesis genes repressed in these datasets were likewise repressed in SLAM cells from IL-1 exposed mice (i.e., enriched in-IL-1 SLAM cells) (**Fig. 3A**). We also identified a common signature of cell cycle and protein synthesis genes, including *Myc* itself, repressed in all three RNA-seq datasets (**Fig. 3B, Table S3**).

**Figure 3.**
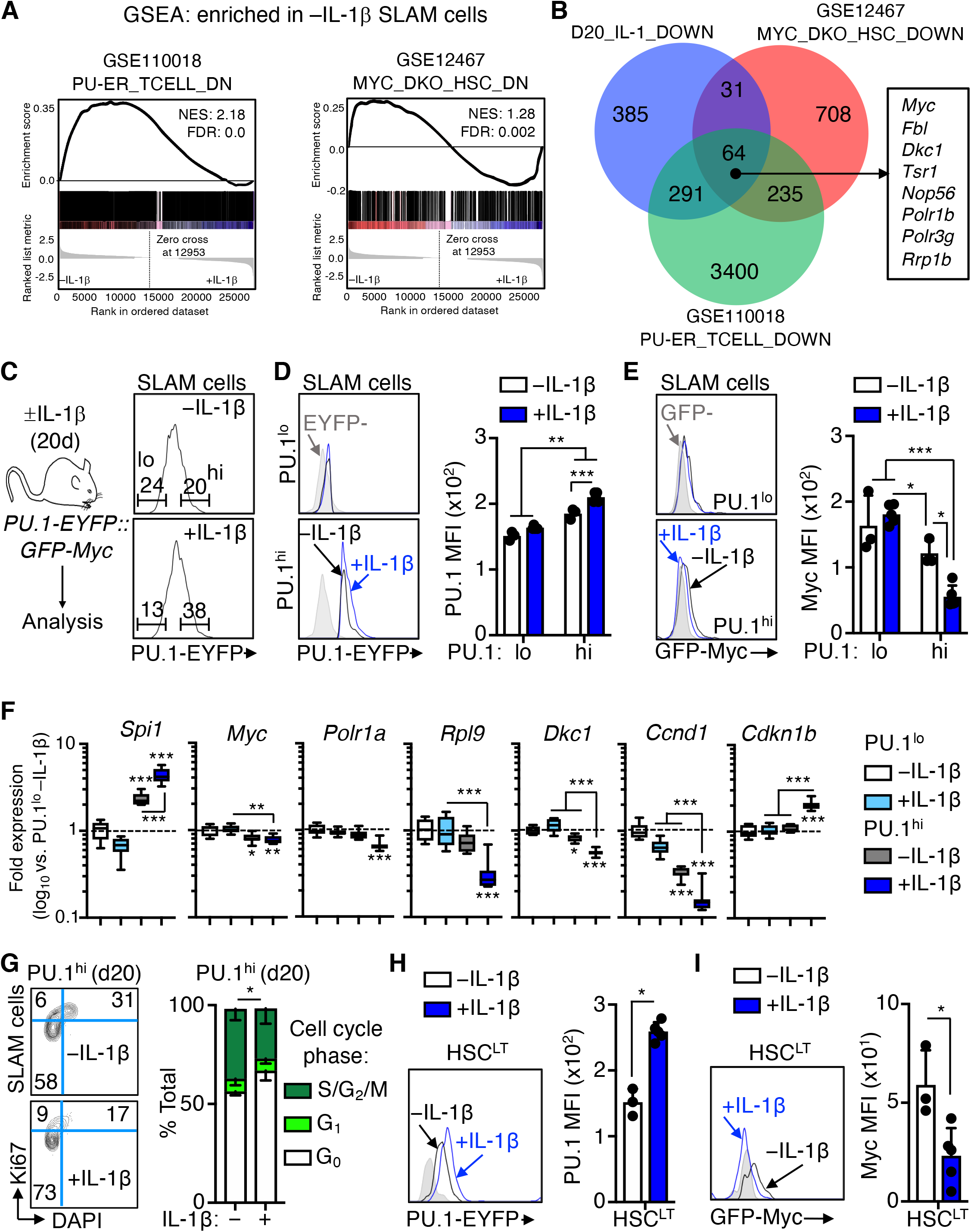
IL-1 induced gene repression is associated with PU.1 activity. **A**) GSEA enrichment of significantly downregulated genes in publicly available datasets versus RNA-seq analysis of SLAM cells from mice treated ± IL-1β for 20d. Data show downregulated genes as negatively enriched in SLAM cells from mice treated 20d +IL-1. See also Table S5. **B**) Venn diagram showing intersections between gene sets in A). A partial list of common genes is depicted at right of the diagram. See also Table S6 for complete list of genes. **C**) Experimental design for analysis of *PU.1-EYFP::GFP-Myc* mice treated ± IL-1β for 20d (left). Representative FACS plots (right) showing gating strategy to identify PU.1^lo^ and PU.1^hi^ SLAM cells in based on PU.1-EYFP expression levels in these mice. **D**) Representative FACS plots (left) and quantification (right) showing PU.1-EYFP expression levels in PU.1^lo^ and PU.1^hi^ SLAM cell fractions from C) (n = 3-IL-1β, 5 +IL-1β). PU.1-EYFP negative control is shown in grey. Individual values are shown with bars representing mean values. Data are representative of two independent experiments. **E**) Representative FACS plots (left) and quantification (right) showing GFP-Myc expression levels in PU.1^lo^ and PU.1^hi^ SLAM cell fractions from C) (n = 3-IL-1β, 5 +IL-1β). GFP-Myc negative control is shown in grey. Individual values are shown with bars representing mean values. Data are representative of two independent experiments. **F**) Quantification by Fluidigm qRT-PCR array of cell cycle and protein synthesis gene expression in PU.1^hi^ and PU.1^lo^ SLAM cells from mice treated ± IL-1β for 20d (n = 8/group). Data are expressed as log_10_ fold expression versus-IL-1β. Box represents upper and lower quartiles with line representing median value. Whiskers represent minimum and maximum values. Data are representative of two independent experiments. **G**) Representative FACS plots (left) and quantification (right) of cell cycle distribution in PU.1^hi^ SLAM cells from mice treated ± IL-1β for 20d (n = 3/group) using Ki67 and DAPI. Data are compiled from two independent experiments. **H**) Representative FACS plots (left) and quantification (right) showing PU.1-EYFP expression levels in HSC^LT^ from mice in C) (n = 3-IL-1β, 5 +IL-1 β). PU.1-EYFP negative control is shown in grey in the FACS plots. Individual values are shown with bars representing mean values. Data are representative of two independent experiments. **I**) Representative FACS plots (left) and quantification (right) showing GFP-Myc expression levels in PU.1^lo^ and PU.1^hi^ SLAM cell fractions from mice in C). GFP-Myc negative control is shown in grey in the FACS plots. Individual values are shown with bars representing mean values. Data are representative of two independent experiments. * p< 0.05, ** p< 0.01, ***p< 0.001 by Mann-Whitney *u*-test or one-way ANOVA with Tukey’s test in F). Error bars represent S.D. See also Figure S3.

We then generated *PU.1-EYFP::GFP-Myc* dual reporter mice using previously published PU.1-EYFP and GFP-Myc reporter mouse strains (Hoppe et al., 2016; Huang et al., 2008) (**Fig. 3C**). As these reporters consist of fluorescent fusion proteins knocked into the endogenous loci, we could directly read out in the same cells the impact of PU.1 on Myc expression. We observed lineage-specific expression patterns of PU.1 and Myc in HSPC from these mice, with granulocyte-macrophage progenitors (GMP; LK/CD41^-^/FcgR^+^; (Pronk et al., 2007)) expressing high levels of PU.1, whereas phenotypic CFU-erythroid (CFU-E; LK/CD41^-^/FcgR^-^/CD150^-^/CD105^+^) and megakaryocyte progenitors (MkP; LK/CD41^+^) expressed high Myc levels (**Fig. S3A-B**). Consistent with our prior work, chronic IL-1 increased PU.1 expression and the frequency of PU.1^hi^ SLAM cells (**Fig. 3C-D**) (Pietras et al., 2016). Notably, PU.1^hi^ SLAM cells had significantly lower GFP-Myc levels than PU.1^lo^ SLAM cells, and Myc levels decreased further following chronic IL-1 exposure (**Fig. 3E**), supporting a model in which IL-1-induced PU.1 represses *Myc*. To assess whether increased PU.1 levels associated with repression of other genes identified by our RNA-seq study, we performed Fluidigm qRT-PCR analysis on PU.1^hi^ and PU.1^lo^ SLAM cells isolated from mice treated ± IL-1 for 20d. Expectedly, *PU.1/Spi1* was further upregulated in PU.1^hi^ SLAM cells by IL-1, whereas *Myc* was repressed (**Fig. 3F**). Importantly, expression of several cell cycle and protein synthesis genes repressed by IL-1 was specifically downregulated in PU.1^hi^ SLAM cells following chronic IL-1 exposure, suggesting PU.1 may repress these genes (**Fig. 3F**). Conversely, IL-1-induced genes such as *Itgam, Flt3*, and *Il1r1* were elevated in PU.1^hi^ SLAM cells and further upregulated by IL-1 treatment (**Fig. S3C**). Consistent with PU.1-mediated repression of cell cycle and protein synthesis genes, *in vivo* chronic IL-1 exposure decreased cell cycle activity in PU.1^hi^ SLAM cells, whereas IL-1 triggered increased cell cycle activity in the PU.1^lo^ SLAM fraction (**Fig. 3G, Fig. S3D**). To link these findings back to phenotypic HSC^LT^ we assessed PU.1-EYFP and Myc-GFP levels in this fraction. As expected, PU.1-EYFP levels were upregulated by IL-1 in phenotypic HSC^LT^, whereas GFP-Myc levels were reduced (**Fig. 3H-I**). We therefore assessed whether PU.1 expression was sufficient to delay HSC^LT^ cell division by analyzing the division kinetics of transgenic *PU.1-ERT* HSC^LT^, which overexpress PU.1 following transgene induction with tamoxifen (4-OHT) (**Fig. S3E**) (Fukuchi et al., 2008). PU.1 overexpression enforced a delay in HSC^LT^ division kinetics consistent with effects of IL-1 (**Fig. S3F-G**). Altogether, our data show that PU.1 is likely the downstream driver of IL-1-induced repression of cell cycle and protein synthesis genes in HSC^LT^.

### PU.1 directly binds cell cycle and protein synthesis genes repressed by IL-1

Given the association identified in our data between high PU.1 levels and IL-1-induced cell cycle restriction in HSC^LT^, we asked whether PU.1 directly interacts with genes repressed by IL-1. We therefore analyzed genome-wide PU.1 binding using ChIP-seq analysis of HSC-enriched LSK/Flk2-/CD150^+^ cells (**Fig. 4A**). Relative to whole cell extract (WCE) controls, we identified 52,127 unique and specific PU.1 peaks throughout the genome, with significant enrichment at 5’-GGAA-3’-containing consensus motifs within 200bp of the peak sites (**Fig. 4B-C** and **Table S4**). Notably, the vast majority of IL-1 DEGs (586/694 IL-1 up; 599/771 IL-1 down) had PU.1 peaks located predominantly in intronic or intergenic regions within ± 20 kb of the gene transcription start site (TSS), likewise within 200bp of 5’-GGAA-3’-containing consensus motifs (**Fig. 4D-F; Fig. S4A-C, Table S4**). GO analysis of IL-1-repressed genes associated with PU.1 ChIP-seq peaks revealed expected enrichment for *rRNA processing, cell proliferation* and *translation* categories (**Fig. 4G and Table S5**). We next analyzed PU.1 peaks associated with the *Myc* gene and found several located within ± 20 kb of the TSS itself, including a peak located within the gene body near the junction of intron 2/exon 3 (**Fig. 4H**) and several located in the 3’ intergenic space (**Table S4**). The intronic Myc peak was also present in two independent published macrophage datasets ((Heinz et al., 2010); GSE21512); indeed, PU.1 peaks identified in our ChIP-seq dataset corresponded closely with peaks in these two datasets (**Fig. S4H, Table S4**). Conversely, IL-1-induced genes with PU.1 peaks were enriched for *cell adhesion, immune response* and other expected gene categories (**Fig. S4D, Table S5**) including the PU.1 target gene *Itgam*, which contained several peaks located both in the gene body as well as in the 5’ intergenic region, including a site near the TSS/promoter as previously characterized and a major peak that may represent an enhancer site (**Fig. S4E**). Interestingly, we also noted that a subset of IL-1-downregulated genes with PU.1 were also significantly enriched for Myc motifs within 1 kb of their TSS relative to upregulated or unchanged genes (**Fig. 4I, Table S4**), further underscoring the negative regulatory relationship between PU.1 and Myc target genes. To further characterize the dynamics of PU.1 interactions with IL-1 target genes we reanalyzed published ChIP-seq data ((Heinz et al., 2010); GSE21512) generated in *PU.1*^-/-^ fetal liver hematopoietic progenitor cells expressing a tamoxifen (4-OHT)-inducible *PU.1* transgene (PU-ER cells; **Fig. S4F**) at different timepoints post induction (Heinz et al., 2010). These cells constitutively express a low level of PU.1 from the transgene, with 4-OHT rapidly inducing active PU.1 protein expression (Walsh et al., 2002). Indeed, a small fraction (177 total) IL-1-repressed genes were constitutively bound by low levels of PU.1 present in the absence of 4-OHT (0h); interestingly those genes were unrelated to cell cycle or protein synthesis (**Fig. 4J-K**). The number of IL-1-repressed genes bound by PU.1 increased significantly and rapidly with 4-OHT treatment, with top GO categories of these genes now including *ribosome biogenesis* and *rRNA processing* (**Fig. 3I-L**). We observed a similar pattern of expression for IL-1-upregulated genes, though in this case *inflammatory response* genes were already bound by PU.1 at 0h (**Fig. 4I-J, Table S5**) and transgene-inducible peaks centered around *cell adhesion* and *mitotic cell division* gene categories (**Fig. S4I-J**) with the latter centered around genes required for late cell cycle stages rather than quiescence exit. Taken together, these data support a model in which PU.1 represses gene expression by directly binding a broad set of cell cycle and protein synthesis genes. They also suggest that at reduced levels, PU.1 does not bind the majority of IL-1-repressed cell cycle and protein synthesis genes, which could ‘prime’ them for aberrant expression in contexts where PU.1 expression and/or function is inhibited.

**Figure 4.**
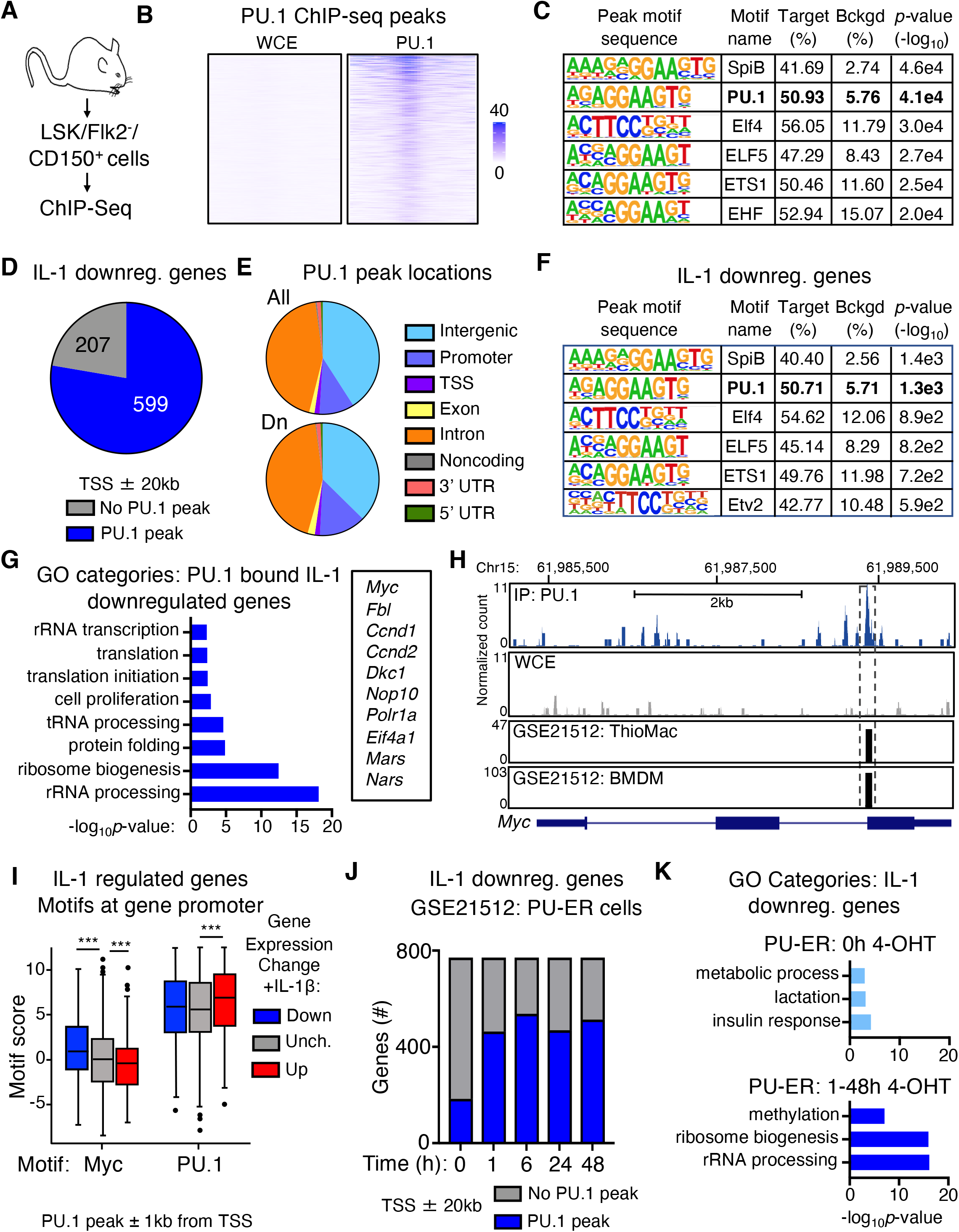
PU.1 directly binds cell cycle and protein synthesis genes repressed by IL-1. **A**) Experimental design for ChIP-seq experiment (n = 2/group). **B**) Heatmap showing PU.1 ChIP-seq peak intensities versus whole cell extract (WCE) control at transcription start site (TSS) ± 1 kb. See also Table S7. **C**) Transcription factor binding site motif enrichment at ChIP-seq peak sites. See also Table S7. **D**) Pie chart comparing IL-1 downregulated genes identified in SLAM cells by RNA-seq analysis in Figure 1 and presence of PU.1 peaks at or near these genes (TSS ± 20 kb). See also Table S7. **E**) Pie chart showing PU.1 peak locations in IL-1 downregulated genes. UTR: untranslated region. **F**) Transcription factor binding site motif enrichment at PU.1 ChIP-seq peak sites in IL-1-downregulated genes (TSS ± 20 kb). **G**) GO category enrichment of IL-1 downregulated DEGs containing PU.1 peaks in C). Representative genes from the indicated categories are shown to the right. Data are expressed as −log_10_ *p*-value. See also Table S8. **H**) UCSC genome browser rendering of a PU.1 peak location in *Myc* gene body. Tracks show PU.1 ChIP-Seq, WCE control, with corresponding peak locations and intensities in thyoglycollate-elicited primary mouse macrophage (ThioMac) and bone marrow-derived macrophage (BMDM) PU.1 ChIP-Seq datasets from GSE21512. **I**) Myc and PU.1 motif enrichment (motif score) at TSS ± 1 kb in IL-1 downregulated, upregulated or unchanged genes containing PU.1 peaks. Box shows upper and lower quartiles with line showing median value; whiskers upper and lower 10^th^ percentile and individual dots represent outliers. ***p ≤ 0.001 based on Wilcoxon rank sum test. **J**) Comparison of IL-1 downregulated genes in SLAM HSC with genes containing PU.1 peaks within TSS ± 20 kb in PU-ER cells ± 4-OHT. Based on PU.1 ChIP-seq datasets in GSE21512. **K**) GO category enrichment of IL-1 downregulated DEGs containing PU.1 peaks in PU-ER cell ChIP-seq dataset at 0h +4-OHT versus combined 1-48h +4-OHT. Top three GO categories are shown. Data are expressed as −log_10_ *p*-value. See also Figure S4.

### IL-1 triggers aberrant protein synthesis and cell cycle activity in PU.1-deficient HSC^LT^

To further explore the impact of reduced PU.1 levels on HSC^LT^ function, we analyzed *PU.1^KI/KI^* mice, which have an ~30% reduction of PU.1 levels in SLAM cells due to a deactivating point mutation knocked into the 14kb upstream *Spi1* autoregulatory binding motif (Staber et al., 2013). As SLAM cells from these mice exhibit de-repression of cell cycle genes including *Ccnd1, E2f* and *Cdk1*, we first compared our RNA-seq data to published gene expression microarray data from *PU.1^+/+^* and *PU.1^KI/KI^* SLAM cells ((Staber et al., 2013); GSE33031). Expectedly, several cell cycle and protein synthesis genes repressed by IL-1, including *Myc*, were significantly upregulated in *PU.1^KI/KI^* SLAM cells (**Fig. S5A**). We therefore assessed gene expression in HSC^LT^ from *PU.1^KI/KI^* mice and *PU.1*^+/+^ littermate controls treated ± IL-1 for 20d by Fluidigm qRT-PCR array (**Fig. 5A**). Hierarchical clustering analysis revealed distinct gene expression changes related to IL-1 exposure PU.1 deficiency (**Fig. S5B**). We confirmed significant reduction of *Spi1* and *Itgam* gene expression in *PU.1^KI/KI^* HSC^LT^ while expression of other IL-1-induced genes was modestly affected (**Fig. S5C-D**). Notably, we observed a pattern of gene expression in which *Cdk1, E2f1* and *Myc* were de-repressed in *PU.1^KI/KI^* HSC^LT^, consistent with prior characterizations of PU.1-deficient HSC (**Fig. 5B-C**) (Rosenbauer et al., 2004; Staber et al., 2013). Increased expression in *PU.1^KI/KI^* HSC^LT^ was also evident for several other cell cycle and protein synthesis genes including *Ccne1, Ccna2* and *Mki67*, whereas IL-1-mediated repression of target genes such as *Rpl9, Eif3a* and *Fbl* was attenuated in *PU. 1^KI/KI^* HSC^LT^ (**Fig. 5B-C**). Chronic IL-1 exposure further enhanced expression of several of these genes including *Myc, E2f1, Ccna1* and *Mki67* in *PU.1^KI/KI^* HSC^LT^, suggesting increased PU.1 levels are required to maintain homeostatic expression of these targets under inflammatory stress (**Fig. 5B-C**). Consistent with our qRT-PCR data, IL-1-exposed *PU.1^KI/KI^* HSC^LT^ expressed higher relative levels of Myc than *PU.1*^+/+^ HSC^LT^ (**Fig. 5D**). Chronic IL-1 also triggered exuberant protein synthesis activity in *PU.1^KI/KI^* HSC^LT^ (**Fig. 5E**). To assess the functional consequences, we assessed cell cycle activity in *PU.1*^+/+^ and *PU.1^KI/KI^* HSC^LT^. Strikingly, IL-1 triggered increased cell cycle activity in *PU.1^KI/KI^* HSC^LT^ (**Fig. 5F**). We independently confirmed IL-1-dependent activation of aberrant protein synthesis and cell cycle activity in HSC^LT^ from conditional *SCL-Cre-ER::PU.1^Δ/Δ^* mice treated ± IL-1 for 20d (**Fig. S5E-G**). In addition, we found that aberrant *PU.1^KI/KI^* HSC^LT^ cell cycle activity and Myc expression could be triggered by a single IL-1 injection, suggesting PU.1 is required to maintain HSC^LT^ quiescence in response to acute inflammatory stimulation as well (**Fig. S5H-J**). Given IL-1 triggered increased protein synthesis and cell cycle activity in PU.1-deficient HSC^LT^, we reasoned this could lead to aberrant expansion of the phenotypic HSC^LT^ pool. Hence, we assessed the number of HSC^LT^ in the BM of *PU. 1*^+/+^ and *PU. 1^KI/KI^* mice treated ± IL-1 for 20d. As anticipated, IL-1 triggered aberrant expansion of phenotypic HSC^LT^ exclusively in the BM of *PU.1^KI/KI^* mice (**Fig. 5G**, **Fig. S6A**). Notably, this phenotype was not confined to the BM, as we also observed expansion of phenotypic SLAM cells exclusively in the spleens of IL-1-treated *PU.1^KI/KI^* mice (**Fig. S6B-C**) Taken together, these data demonstrate that aberrant protein synthesis and cell cycle activity are emergent properties of PU.1-deficient HSC^LT^ that can be triggered by IL-1 and result in expansion of phenotypic HSC^LT^ in the BM and extramedullary sites (**Fig. 5H**) reminiscent of a pre-leukemic and/or clonal hematopoiesis-like state.

**Figure 5.**
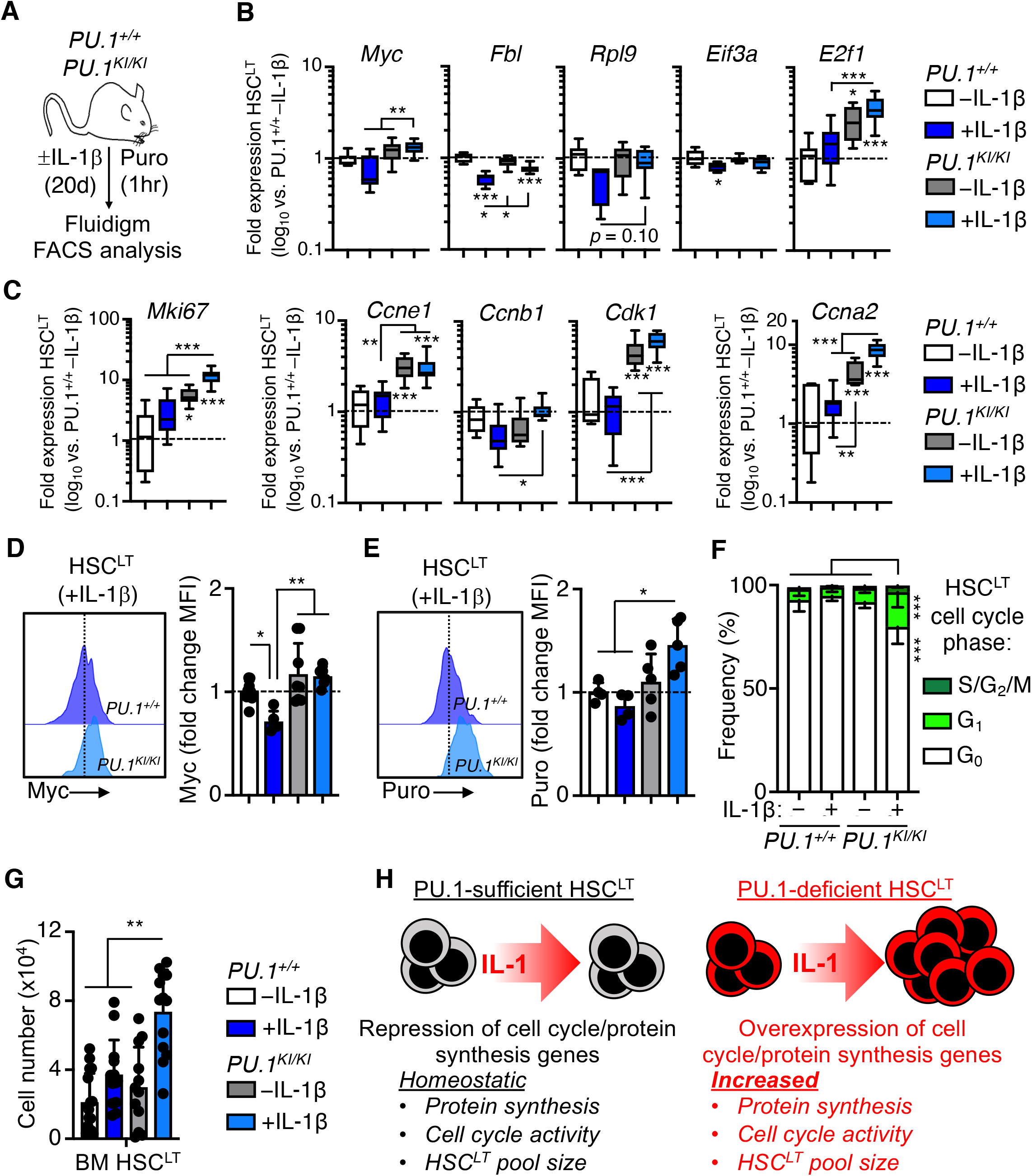
IL-1 triggers aberrant protein synthesis and cell cycle activity in PU.1-deficient HSC^LT^. **A**) Experimental design for analyses of HSC^LT^ from *PU. 1^+/+^* and *PU. 1^KI/KI^* mice treated ± IL-1β for 20d. **B**) Quantification by Fluidigm qRT-PCR array of protein synthesis gene expression in HSC^LT^ from A). Data are expressed as log_10_ fold expression versus-IL-1β. Box represents upper and lower quartiles with line representing median value. Whiskers represent minimum and maximum values. **C**) Quantification by Fluidigm qRT-PCR array of cell cycle gene expression in HSC^LT^ from A). Data are expressed as log^10^ fold expression versus-IL-1β. Box represents upper and lower quartiles with line representing median value. Whiskers represent minimum and maximum values. **D**) Intracellular flow cytometry analysis of Myc protein levels in HSC^LT^ from *PU.1^+/+^* and *PU.1^KI/KI^* mice treated ± IL-1β for 20d (n = 4-9/group) Data are expressed as fold change of mean fluorescence intensity (MFI) versus -IL-1 β. Individual values are shown with bars representing mean values. Data are compiled from two independent experiments. **E**) Intracellular flow cytometry analysis of puromycin (Puro) incorporation in HSC^LT^ from *PU.1^+/+^* and *PU.1^KI/KI^* mice treated ± IL-1β for 20d (n = 4 *PU.1^+/+^*, 5 *PU.1^KI/KI^).* Data are expressed as fold change of mean fluorescence intensity (MFI) versus-IL-1β. Individual values are shown with bars representing mean values. Data are from one experiment. **F**) Quantification of cell cycle phase distribution in HSC^LT^ from mice in E). Data are compiled from two independent experiments. **G**) Quantification of BM HSC^LT^ from mice in A). Individual values are shown with bars representing mean values. * p< 0.05, ** p< 0.01, ***p< 0.001 by one-way ANOVA with Tukey’s test. Error bars represent S.D. See also Figure S5 and Figure S6.

## Discussion

Here, we show that chronic IL-1 exposure represses a broad set of genes regulating HSC^LT^ cell cycle and protein synthesis activity. We show this gene program is linked to restricted HSC^LT^ cell cycle entry and thus underwrites the quiescent phenotype of HSC under chronic inflammatory stress. Notably, PU.1 is a crucial driver of this program, with PU.1 itself binding the majority of target genes repressed by IL-1. Strikingly, our data demonstrate this molecular state leaves PU.1-deficient HSC^LT^ poised to exit quiescence and support aberrant expansion of the phenotypic HSC^LT^ pool when triggered by IL-1. Our data identify a PU.1-driven molecular mechanism enforcing HSC quiescence under chronic inflammatory stress and support a model in which PU.1 expression serves as a critical limiting mechanism that regulates HSC cell cycle activity and pool size in this context.

Tight regulation of cell cycle activity is crucial for maintaining the long-term functional integrity of the HSC pool (Matsumoto et al., 2011; Pietras et al., 2011). This is particularly true under inflammatory stress conditions, where increased proliferative activity can lead to overproduction of blood cells, rapid HSC attrition and BM failure (Essers et al., 2009; King et al., 2011; Matatall et al., 2016; Pietras, 2017; Pietras et al., 2014; Takizawa et al., 2017). Highly enriched HSC fractions can maintain a quiescent phenotype and long-term engraftment capacity even under a variety of inflammatory stress conditions (Bujanover et al., 2018; Hernandez et al., 2020; Pietras et al., 2014; Pietras et al., 2016; Rabe et al., 2020; Wilson et al., 2008; Zhao et al., 2019), implying the existence of mechanism(s) that prevent spurious HSC cell cycle entry.

We previously reported that IL-1 activates precocious myeloid differentiation in HSC (Pietras et al., 2016). Here, we demonstrate that IL-1 activates a PU.1 program that restricts HSC cell cycle activity and myeloid expansion. Here, we show that IL-1/PU.1 enforces HSC quiescence by initiating the repression of a broad set of cell cycle and protein synthesis genes. We made similar observations in HSC from mice with collagen-induced arthritis (CIA) (Hernandez et al., 2020), suggesting repression of cell cycle and protein synthesis genes by PU.1 is a common mechanism triggered in HSC by at least a subset of inflammatory stimuli. Indeed, it is likely that signaling through at least some TLRs, which share the downstream MyD88 signaling adapter with IL-1, may also directly trigger this mechanism. On the other hand, analysis of HSC from PU.1-EYFP mice treated acutely with TLR3 and −4 ligands poly I:C and LPS respectively, show that PU.1 is induced via a TNF-dependent mechanism (Etzrodt et al., 2019; Yamashita and Passegue, 2019). Notably, TLR4-dependent HSC proliferation was shown to be triggered by TRIF rather than MyD88 (Takizawa et al., 2017), consistent with our finding that IL-1 itself does not robustly activate HSC^LT^ proliferation. On the other hand, cytokines such as IFNs do not robustly activate PU.1 in a direct fashion (Etzrodt et al., 2019), suggesting other mechanism(s) may enforce HSC quiescence under chronic IFN stimulation, such as activation of p53 and/or the mRNA translation-blocking activity of several interferon-stimulated genes (ISGs) (Li et al., 2015; Pietras et al., 2014).

The activating versus suppressive functions of the IL-1/PU.1 axis may at first appear paradoxical. Indeed, previous reports indicate that acute IL-1 triggers HSC cell cycle activity (Hemmati et al., 2019; Pietras et al., 2016; Weisser et al., 2016). Here, we show that IL-1 does not trigger cell cycle activity in PU.1-sufficient HSC^LT^, even in the context of acute stimulation. Importantly, these studies used the more general SLAM definition to enrich for phenotypic HSC. Here, we show that EPCR^+^/7CD34^-^ HSC^LT^ remain largely quiescent following acute IL-1 stimulation, whereas IL-1 robustly activates proliferation of ECPR^-^ fractions within the SLAM gate that can contribute to ‘emergency’ hematopoiesis. These findings suggest that a sizeable fraction of HSC^LT^ remain largely dormant in response to inflammatory stressors. This is in line with previous work showing that poly I:C specifically triggers proliferation of CD41^hi^ SLAM cells that constitute a pre-megakaryocyte lineage (Haas et al., 2015) and that a subset of CD49b^lo^ HSC, which overlaps phenotypically with the EPCR^+^7CD34^-^ HSC^LT^ fraction used here (Rabe et al., 2020), can remain dormant even following 5-FU myeloablation (Zhao et al., 2019). Likewise, BrdU label-retention experiments show that while stressors such as LPS can trigger proliferation in HSC-enriched cells, a sizeable fraction of these cells retains the BrdU label (Takizawa et al., 2017; Wilson et al., 2008) suggesting mechanism(s) remain in place to limit their proliferative activity.

PU.1 has previously been characterized as a negative regulator of cell proliferation (Delestre et al., 2017; Fukuchi et al., 2008; Oikawa et al., 1999; Solomon et al., 2017; Ziliotto et al., 2014). In the context of quiescent HSC^LT^, our data show that increased PU.1 activity serves as more as a rheostat than a bona fide proliferation block, erecting enough of an activation barrier to prevent spurious proliferation during inflammatory challenge while still ‘priming’ HSC for myeloid differentiation as evidenced by simultaneous upregulation of myeloid determinant genes in IL-1-exposed HSC^LT^. The connection between cell cycle inhibition and myeloid differentiation likely centers on cell cycle lengthening as a mechanism to promote PU.1 accumulation and myeloid differentiation in hematopoietic progenitors (Kueh et al., 2013). This is a function common to numerous myeloid transcription factors, including *Cebpa* family members and Gfi1, which is itself a PU.1 target (Hock et al., 2004; Porse et al., 2005; Pulikkan et al., 2017; Staber et al., 2013; Umek et al., 1991; Zeng et al., 2004). Thus, induction of a PU.1-dependent cell cycle restriction gene program in HSC^LT^ under inflammatory stress conditions is consistent with a model in which PU.1 promotes efficient generation of myeloid progenitors to reconstitute hematopoiesis, while protecting the HSC compartment from damage and/or depletion. Absence of this program likely underlies the failure of *PU.1^KI/KI^* mice to reconstitute hematopoiesis following 5-FU treatment, which we interpret as a coupled effect of increased HSC proliferation leading to apoptosis, and inefficient generation of new myeloid progenitors due to reduced PU.1 accumulation. Altogether, this mechanism may be a remarkable example of parsimony in a biological system, wherein a single program can mediate distinct functional outcomes based on cell type and context.

Our data agree with prior studies showing that PU.1 deficiency leads to de-repression of *Myc* and numerous cell cycle genes including *Cdk1* and *E2f1* (Rosenbauer et al., 2004; Staber et al., 2013; Will et al., 2015), which is exacerbated by IL-1 exposure. We also find that inhibition of protein synthesis in cultured HSC^LT^ using omacetaxine phenocopies the delayed cell cycle entry observed with IL-1 stimulation, suggesting that PU.1 inhibition of protein synthesis genes contributes to enforced HSC quiescence. Notably, Oma is an FDA approved therapy for chronic myelogenous leukemia and can directly ablate MDS stem cells with aberrant levels of protein synthesis (Stevens et al., 2018). Protein synthesis and cell cycle activity are closely linked, as cell cycle entry and mitosis require a significant amount of new protein. Hence in eukaryotic cells, deletion and/or knockdown of key genes regulating protein synthesis such as translation elongation initation factors (eIFs), several of which are repressed by IL-1 in our dataset, results in delayed cell cycle entry (Polymenis and Aramayo, 2015). The importance of eIFs in initiating HSC cell cycle activity has been illustrated by elegant work in which dual deletion of *4E-BP1/2* (which negatively regulate translation by inhibiting eIF4E) leads to increased protein synthesis, aberrant cell cycle activity and HSC expansion (Signer et al., 2016). Likewise, deletion of *Pten*, which dephosphorylates and inhibits the eIF activator mTOR, leads to similar aberrant HSC activity and hypersensitivity to inflammatory factors such as G-CSF and IFNs (Porter et al., 2016; Signer et al., 2014). This phenotype is strikingly reminiscent of IL-1-exposed PU.1-deficient HSC^LT^ in our model. In a similar line, inflammatory initiation of a cell cycle primed ‘G_alert_’ state in HSC and other cells requires activation of mTOR (Rodgers et al., 2014). We also identified decreased expression of ribosomal RNA genes, as well as genes such as *Fbl* required to process them for assembly into ribosomes. While cells synthesize ribosomes independently of cell cycle phase, inhibition of rRNA production can halt cell cycle activity (Polymenis and Aramayo, 2015) with accompanying decreases in cell size, similar to our findings in HSC^LT^ exposed to IL-1 and/or Omacetaxine in culture. Along these lines, HSC carrying a hypomorphic allele of the ribosomal protein *Rpl24* exhibit decreased proliferative activity and can rescue the phenotype of *Pten*-deficient HSC (Signer et al., 2014). Likewise, IL-1 downregulates expression of *c-Myc*, a crucial upstream regulator of protein synthesis and cell cycle genes that leads to defective cell cycle entry if deleted in stem cells (Laurenti et al., 2008; Scognamiglio et al., 2016; Wilson et al., 2004). Together these data suggest that fine tuning of protein synthesis is essential for maintaining HSC quiescence and function (Signer et al., 2014). Interestingly, despite IL-1-mediated downregulation of protein synthesis genes and slowed cell cycle progression in our *in vitro* cultures, we do not observe decreased protein synthesis rates in IL-1-exposed HSC^LT^ *in vivo*. Repression of protein synthesis genes may therefore dampen the effects of IL-1-driven mitogenic signaling and maintain homeostatic protein synthesis activity rather than block it altogether, which *in vitro* is read out as slowed cell cycle progression in response to the stress of being placed in culture. IL-1 activates numerous signaling pathways that directly impact protein synthesis and cell cycle activity including Akt, p38 MAPK, MEK/ERK and RAS (Dinarello, 2018). Further studies can identify which pathway(s) are potential triggers that activate aberrant protein synthesis and cell cycle activity in PU.1-deficient HSC, and the extent to which inhibiting them can rescue this phenotype.

Dysregulated inflammatory signaling has emerged as a key partner in the development and/or progression of hematological malignancy (Barreyro et al., 2018; Pietras, 2017). Aberrant signaling via IL-1 and the downstream IRAK/TRAF pathway shared by IL-1R and Toll-like receptors has been implicated as a driver of survival and/or expansion of MDS and myeloid leukemia stem cells (LSC) (Agerstam et al., 2016; Askmyr et al., 2013; Barreyro et al., 2012; Carey et al., 2017; Muto et al., 2020; Smith et al., 2019; Zhang et al., 2016). Indeed, blockade of these pathways can limit and/or reverse disease progression (Carey et al., 2017; Mitchell et al., 2018; Rhyasen et al., 2013; Zhang et al., 2016). However, the mechanism(s) by which inflammatory signaling initiates expansion of oncogenically mutated HSC remain obscure. Chronic inflammation, often associated with increased IL-1 activity, is a common phenotype in individuals at risk for hematological malignancy, which can be related to several pre-existing factors including aging/tissue decline, genotoxin exposure, metabolic dysfunction and/or autoimmune disease (Barreyro et al., 2018; Pietras, 2017). Our data suggest that while chronic IL-1 exposure can induce expansion of myeloid-biased hematopoietic progenitors and mature myeloid cells, this phenotype is essentially self-limiting and does not deterministically progress to malignancy. These data are consistent with the overall rarity of hematological malignancy amongst individuals with chronic inflammatory phenotypes, despite the relative increase in risk (Anderson et al., 2009; Ganan-Gomez et al., 2015). On the other hand, loss-of function mutations in *PU.1* itself are rarely observed in hematological malignancy, though a wide array of myeloid leukemia-associated oncogenic lesions can interfere with PU.1 expression or function, including *PML/RARA, AML1-ETO, NPM1c*, and mitogenic kinase mutations in *BCR/ABL* and *Flt3^ITD^* (Gerloff et al., 2015; McKenzie et al., 2019; Mueller et al., 2006; Noguera et al., 2016; Vangala et al., 2003; Yang et al., 2012). In addition, *TET2* and/or *DNMT3A* mutations associated with early oncogenesis may interfere with PU.1 function due to aberrant methylation of PU.1 binding sites (Kaasinen et al., 2019). Hence, PU.1 deficiency related to indirect effects of mutations in other genes may serve as a crucial ‘nexus’ that primes HSC to activate aberrant protein synthesis and cell cycle activity when triggered by inflammatory signals associated with bone marrow pathogenesis like IL-1 and TNF. These findings are consistent with the theory of Adaptive Oncogenesis, which stipulates that environmental factors associated with tissue decline, such as inflammation, are crucial drivers of oncogenesis by selecting for emergent phenotypes such as increased proliferation or survival that are linked cancer-associated mutations. Hence, our data support a model of leukemogenesis in which oncogenic mutations and inflammatory signals collaborate to activate emergent proliferative phenotypes in stem cells, that in turn lead to aberrant expansion of downstream progenitors and myeloid cells that can initiate disease. They also provide rationale for exploring the use of anti-inflammatory therapies as a means of preventing and/or delaying leukemogenesis in at-risk individuals.

## Acknowledgements

We thank Garrett Hedlund for expert assistance with flow cytometry resources. This work was supported by R01 DK119394, K01 DK098315, the Boettcher Webb-Waring Early Career Investigator Award and the Cleo Meador and George Ryland Scott Endowed Chair in Hematology (to E.M.P.), R01 AG067584 (to J.D.), F31 HL138754 (to J.L.R.), F30 CA210383 and T32 AG000279 (to K.C.H.), Swiss National Science Foundation grant 179490 to T.S, and the National Science Foundation Graduate Research Fellowship Program (NSF GFRP; to T.S.M.). This work was supported in part by the University of Colorado Cancer Center Flow Cytometry Shared Resource, funded by NCI grant P30 CA046934.

## Author Contributions

Conceptualization: E.M.P. Methodology: E.M.P., J.D., D.L., K.C.H. and T.S.; Formal Analysis: D.L., N.A., H.K, J.R.M. and B.M.S.; Investigation: J.S.C., J.L.R., D.L., G.H., R.L.G. Z.K., T.S.M., K.C.H., N.A, B.M.I. and E.M.P.; Resources: H.N., T.S. and R.S.W.; Writing–Original Draft: E.M.P. and J.S.C.; Writing–Review & Editing: E.M.P., J.S.C. and J.D.; Supervision: E.M.P., J.D., C.T.J., J.A., and T.S.; Funding Acquisition: E.M.P. and J.D.

## Methods

### Mice

Wild-type C57BL/6, congenic *B6.SJL-Ptprc^a^Pepc^b^/BoyJ* (Boy/J) mice, *MyD88^-/-^* mice and *GFP-Myc* mice were obtained from The Jackson Laboratory. *PU.1^KI/KI^* mice and PU.1^*flox*^mice (Staber et al., 2013) were a kind gift of Dr. Dan Tenen (Beth Israel). *PU.1-EYFP* mice (Hoppe et al., 2016) were a kind gift of Dr. Claus Nerlov (MRC Weatherall Institute). *PU.1ERT2* mice were kindly provided by Dr. Hideyaki Nakajima. All animal experiments were conducted in accordance with a protocol approved by the Institutional Animal Care and Use Committee (IACUC) at University of Colorado Anschutz Medical Campus (protocol number 00091).

### *In vivo* procedures

For *in vivo* IL-1 stimulation, IL-1β (Peprotech) was resuspended in sterile D-PBS/0.2% BSA. 0.5 μg of IL-1 or PBS/BSA alone was injected intraperitoneally (i.p.) in a 100 μl bolus once per day for either one or 20 days, as previously described (Pietras et al., 2016; Rabe et al., 2020). *In vivo* puromycin labeling assays were performed as previously described (Chapple et al., 2018), with 500 μg of puromycin (Life Technologies) injected i.p. one hour prior to euthanasia.

### Flow cytometry

Analysis of BM cell populations was performed using a similar protocol as previously described (Pietras et al., 2016; Rabe et al., 2020). Bone marrow was flushed from femurs and tibiae of mice using staining media (SM; Hanks Buffered Saline Solution + 2% FBS) and a syringe equipped with a 21G needle. Cells were subsequently resuspended in 1x ACK (150 mM NH_4_Cl/10 mM KHCO_3_) to remove erythrocytes, washed with SM, filtered through 70μm mesh to remove debris, resuspended in SM and counted on a Vicell automated counter (Beckman Coulter). For quantification of immature hematopoietic stem and progenitor cells, 1×10^7^ BM cells were blocked with purified rat IgG (Sigma-Aldrich) and stained for 30 minutes on ice with SM containing the following antibodies: PE-Cy5-conjugated anti-CD3, CD4, CD5, CD8, Gr-1 and Ter119 as a lineage exclusion stain, Flk2-biotin, CD34-FITC, EPCR-PE, Mac-1-PE/Cy7, CD16/32-APC, CD48-A700, and cKit-APC/Cy7. Cells were subsequently washed, resuspended in a 1:4 dilution of Brilliant Staining Buffer (Becton-Dickenson) in SM containing Sca-1-BV421, CD41-BV510, CD105-BV711, CD150-BV785, and streptavidin (SA)-BV605 and incubated for 30 minutes on ice. For analysis of *GFP-Myc::PU.1-EYFP* cells, a custom 510/20 nm (GFP) and 550/30 nm (EYFP) bandpass filter setup (Becton Dickenson) was used to discriminate the two fluorescent proteins. BM cells from single-positive mice were used for compensation controls and fluorescence minus one (FMO) controls, and GFP-/YFP-BM cells were stained alongside and used as negative controls. For analysis of mature BM cell populations, cells were blocked as above and stained for 30 minutes on ice with the following antibodies: Gr-1-Pacific Blue, Ly6C-BV605, B220-BV786, CD4-FITC, CD8-PE, Mac-1-PE/Cy7, IgM-APC, CD3-A700, and CD19-APC/Cy7. Following surface staining, cells were washed with SM, resuspended in SM containing 1 μg/ml propidium iodide (PI) and immediately analyzed on a 3-laser, 12 channel FACSCelesta analyzer (Becton-Dickenson) or a 5-laser, 18-channel BD Fortessa. Cell cycle analyses were performed as previously described. For analysis of spleen cells, spleens were removed and minced through a 70 μm filter basket (VWR) to make a single-cell suspension. Splenocytes were treated with ACK buffer as above and subsequently processed for staining of mature and immature hematopoietic populations as described above. Cell cycle analysis was performed using protocols similar to previous publications (Hernandez et al., 2020; Jalbert and Pietras, 2018), 1×10^7^ BM cells were stained for 30 min. on ice with the following antibodies: PE-Cy5-conjugated anti-CD3, CD4, CD5, CD8, Gr-1 and Ter119 as a lineage exclusion stain, Flk2-biotin, CD34-FITC, EPCR-PE, Sca-1-PE/Cy7, CD48-A700, and c-Kit-APC/Cy7. Cells were washed with SM and resuspended in 1:4 Brilliant buffer:SM containing SA-BV605 and CD150-BV785, incubated for 30 minutes on ice, washed in SM and fixed for 20 minutes at room temperature (RT) with Cytofix/Cytoperm (Becton-Dickenson). Cells were subsequently washed with 1x PermWash buffer (Becton-Dickenson), permeablized for 10 minutes at RT with Perm Buffer Plus (Becton-Dickenson), washed in PermWash and re-fixed for 5 minutes at RT with CytoFix/Cytoperm. Cells were subsequently washed in PermWash and incubated with anti-Ki67 antibody in PermWash buffer for 1hr at RT. Cells were then washed and either stored at 4°C or resuspended in SM containing 2 μg/ml DAPI and analyzed on a BD LSRII analyzer equipped with a UV laser (Becton-Dickenson). Myc staining was performed as previously described (Freire and Conneely, 2018). 10^7^ BM cells were stained for 30 minutes on ice with SM containing the following antibodies: PE-Cy5-conjugated anti-CD3, CD4, CD5, CD8, Gr-1 and Ter119 as a lineage exclusion stain, Flk2-biotin, CD34-FITC, EPCR-PE, Mac-1-PE/Cy7, CD48-A700, and cKit-APC/Cy7, washed in SM and stained for 30 min in 1:4 Brilliant buffer:SM containing Sca-1-BV421, SA-BV605 and CD150-BV785. Cells were subsequently washed in SM and fixed for 20 minutes with CytoFix/CytoPerm. Cells were washed with PermWash and blocked in PermWash for 1 hr at RT, followed by washing and a 1hr RT incubation with anti-Myc purified antibody. Cells were subsequently stained with an A647-conjugated anti-rabbit Fab for 30 min at RT, washed in PermWash, resuspended in SM and analyzed on an LSRII. Puromycin stainings were performed as previously described (Chapple et al., 2018). Cells were surface stained and fixed using the same approach as for Myc staining. Following fixation, cells were stained with anti-puromycin antibody diluted in PermWash buffer for 1 hr at RT, washed and stained with an anti-mouse IgG2a antibody for 30 min, washed once more and resuspended in SM for analysis on an LSRII. A complete list of FACS antibodies and dilutions used can be found in **Table S6**.

### Cell sorting

For HSC isolation by cell sorting, arm, leg, pelvic bones and spines were isolated from mice as previously described (Pietras et al., 2016; Rabe et al., 2020). Bones were crushed in SM using a mortar and pestle and depleted of erythrocytes using 1x ACK. Cells were subsequently placed atop a Histopaque 1119 gradient (Sigma-Aldrich) and centrifuged. To enrich c-Kit^+^ cells, BM cells were incubated on ice for 20 minutes with c-Kit microbeads (Miltenyi Biotec, 130-091-224; 5 μl/100 μl SM per mouse), washed with SM and enriched on an AutoMACS Pro magnetic cell separator (Miltenyi Biotec). Enriched cells were subsequently washed, blocked with rat IgG and stained for 30 minutes on ice with the following antibodies: PE-Cy5-conjugated anti-CD3, CD4, CD5, CD8, Gr-1 and Ter119 as a lineage exclusion stain, Flk2-biotin, CD34-FITC, ECPR-PE, Mac-1-PE/Cy7, CD48-A700 and c-Kit-APC/Cy7. Cells were subsequently washed with SM and resuspended in a 1:4 dilution of Brilliant Staining Buffer in SM containing Sca-1-BV421, SA-BV605 and CD150-BV785. For B220+ cell isolation by cell sorting, spleens were harvested, pressed over a 70-micron filter and depleted of erythrocytes. To enrich for B220+ cells, splenocytes were incubated on ice for 20 minutes with B220 microbeads (Miltenyi Biotec,; 20 μL/1000 μL SM per mouse), washed with SM and enriched on an AutoMACS Pro. Enriched cells were washed, blocked with rat IgG and stained with B220-APC for 15 minutes on ice. Purified B220+ cells were immediately fixed for 20 minutes at room temperature (100μL, BD Cytofix/Cytoperm), washed with SM and stored at 4°C.

### In vitro culture assays

Purified HSC^LT^ were cultured using a similar protocol as previously published (Pietras et al., 2016). Double-sorted cells were grown in in StemPro34 containing Stempro supplement, Antibacterial-antimycotic (100x, Gibco), L-Glutamine (100x, Gibco), SCF (25 ng/mL), TPO (25 ng/mL), IL-3 (10 ng/mL), GM-CSF (20 ng/ml), Flt3L (50 ng/mL), IL-11 (50 ng/mL), EPO (4 U/mL), and IL-1β (25 ng/mL) and incubated at 37°C, 5% O_2_, 5% CO_2_ for either 12 or 24 hours. Cell culture grade puromycin (1nM, Gibco) was added 1h prior to harvesting cells to monitor protein synthesis rates. In the experiments containing Omacetaxine (100nM, Teva Pharmaceuticals), cells were cultured in the presence of omacetaxine for 1h prior to IL-1β addition and remained in cultures for the duration of the experiment. Following indicated times in culture, cells were harvested and either resorted based on viability for Fluidigm gene expression analysis or were immediately fixed for 20 min at room temperature (100μL, BD cytofix/cytoperm) for Ki67/DAPI analysis or puromycin incorporation assays. Following fix/perm, 5×10^5^ sorted, fixed B220+ spleen cells were added to each sample as carrier cells and excluded during analysis based on B220 expression (Matatall et al., 2018).

### Single-cell tracking analysis

Time-lapse experiments were conducted at 37 °C, 5% O_2_, 5% CO_2_ on μ-slide VI^0,4^ channels slides (IBIDI), coated for 1h at room temperature with 5 or 10 μg/mL anti-CD43-biotin as described (Loeffler et al., 2018). Cells were cultured in phenol red-free IMDM supplemented with 20% BIT, 2 mM L-Glutamine, 50 μM 2-mercaptoethanol and 50 U/mL penicillin, 50 μg/mL Streptomycin, 25 ng/mL SCF, 25 ng/mL TPO, 10ng/mL IL-3, 20ng/ml GM-CSF, 50ng/mL Flt3L, 50ng/mL IL-11 and 4U/mL EPO. 25ng/mL IL1b and / or 500nM tamoxifen (4-OHT, Sigma) was added as indicated. Images were acquired using a Nikon-Ti Eclipse equipped with a linear encoded motorized stage, Orca Flash 4.0 V2 (Hamamatsu), Spectra X fluorescent light source (Lumencor) and The Cube (Life Imaging Service) temperature control system. White light emitted by Spectra X was collimated and used as a transmitted light for bright field illumination via a custom-made motorized mirror controlled by Arduino UNO Rev3 (Arduino). Fluorescent images were acquired using optimized filter sets: eGFP (470/40; 495LP; 525/50), YFP (500/20; 515LP; 535/30), mKO2 (546/10; 560LP; 577/25), mCherry (550/32; 585LP; 605/15), Cy5 (620/60; 660LP; 700/75; all AHF) to detect GFP-cMYC, PU.1-YFP, CD71-PE, TMRM and CD71-APC, respectively. Time intervals of bright field and fluorescent image acquisition were chosen to minimize phototoxicity. Images were acquired using a 10x CFI Plan Apochromat λ objective (NA 0.45). Single-cell tracking and image quantification were performed using self-written software as described (Hilsenbeck et al., 2016; Loeffler et al., 2018). Software used for data acquisition of time-lapse imaging data is published and open-sourced (YouScope v.2.1; http://langmo.github.io/youscope/). Software for single-cell tracking and fluorescence quantification used in this study is published and open-sourced (https://doi.org/10.1038/nbt.3626). Segmentation software is published and open-sourced (https://academic.oup.com/bioinformatics/article/33/13/2020/3045025). Acquired 16-bit images with 2048×2048 pixel resolution were saved as.png and linearly transformed to 8-bit using channel optimized white points and corrected for background and shading (Peng et al., 2017) before analysis. Brightfield images were used for segmentation using fastER (Hilsenbeck et al., 2016), eroded to reduce segmentation artifacts caused by close cell proximity (Settings: Morphological transformation x = 3, y = 3, op = 2, shape: 2) and subsequently dilated (Settings: simple dilation 6) to ensure proper segmentation and quantification results. Tracking and quantification of fluorescence channels were done as described (Hilsenbeck et al., 2016) and analyzed using Matlab 2018b (Mathworks).

### RNA-seq

RNA was isolated from individual pools of 1×10^4^ to 2×10^4^ double-sorted SLAM cells using an RNEasy Micro kit (Qiagen). RNA was quantified and quality checked using an Agilent Bioanalyzer 2100 (Agilent). 1ng of total RNA was preamplified using the SMARTer Ultra Low Input kit v4 (Clontech) according to the manufacturer’s instructions and cDNA quality was assessed using the Qubit Fluorometer (Life Technologies) and the Bioanalyzer. 150pg of cDNA was used to generate Illumina sequencing libraries using the NexteraXT kit (Illumina) according to the manufacturer’s instructions. Libraries were subsequently hybridized to an Illumina single-end flow cell and amplified using the cBot (Illumina). Sequencing of libraries was performed on Illumina HiSeq 2500v4 high-throughput sequencer. Single-ended reads of 100nt were generated for each sample and demultiplexed using bcl2fastq v1.8.4. Trimmomatic v0.32 was used for quality filtering and adapter removal. Processed reads were mapped to the mouse genome GRCm38.p4 (mm10/mg38) using STAR_2.4.2a. Raw read counts were obtained via htseq-count v0.6.1 and Gencode-M12 gene annotations. Differential expression analyses were performed using deSeq2 in Bioconductor. Genes with *p*_adj_ < 0.05 were considered significant differentially expressed genes (DEGs). Upstream Regulator analysis of DEGs was performed using Ingenuity Pathway Analysis software on default settings. GSEA analyses were performed on DEGs using default settings. GO analyses were performed using DAVID v6.8 (david.ncifcrf.gov) (Huang da et al., 2009). Venn diagrams were generated from DEG lists using IteractiVenn (interactivenn.net) (Heberle et al., 2015).

### ChIP-seq

ChIP-seq analysis was performed on LSK/Flk2^-^/CD150^+^ cells sorted from BM of 8 C57BL/6 mice per ChIP. Cells were cross-linked in 0.75% formaldehyde, pelleted and frozen until the ChIP procedure. Pellets were thawed on ice, resuspended in lysis buffer (1% SDS, 10mM EDTA, 50 mM Tris-HCL pH 8.1 + protease inhibitor) for 10 min. Volume was brought up to 1 ml with ChIP dilution buffer (16.7mM Tris-HCl pH 8.1, 167mM NaCl, 0.01% SDS, 1.1% Triton X-100, 1.2mM EDTA + protease inhibitor) and samples were sonicated with a Branson sonicator to shear chromatin. Chromatin was subsequently incubated with 1 μg PU.1 antibody (Santa Cruz; sc-390405) and rotated overnight at 4°C. Next, ChIPs were added to a 50:50 mix of pre-washed Protein A/G Dynabeads (50 μl; Thermo Fisher; 10001D/10009D) and rotated overnight at 4°C. The next day, 175 μl of RIPA/140 mM NaCl buffer (0.1% DOC, 0.1%SDS,1% Triton X-100,140mM NaCl,1mM EDTA, 20mM Tris-HCl pH 8.1) was added to each tube and beads were transferred to a 96-well plate on a magnet, washed twice with cold RIPA/500ml NaCl buffer (0.1% DOC, 0.1%SDS,1% Triton X-100,140mM NaCl,1mM EDTA, 20mM Tris-HCl pH 8.1), twice with cold LiCl buffer (0.25M LiCl,1% NP40,1% Na Deoxycholate, 1mM EDTA,10mM Tris-HCl pH 8.1), and twice with room-temperature TE buffer (10mM Tris-HCl pH8.0, 1mM EDTA pH 8.0). Chromatin was eluted by adding 50 μl elution buffer (10mM Tris-Cl pH 8.0, 5mM EDTA, 300mM NaCl, 0.1% SDS and 5mM DTT) and 8 μl reverse cross-linking buffer 250mM Tris-HClpH 6.5, 1.25M NaCl, 62.5mM EDTA, 5mg/ml Proteinase K, and 62.5ug/ml RNAse A) and incubated at 65°C overnight. ChIP material was quantified on a Qubit spectrometer (Invitrogen) according to manufacturer’s instruction. Library construction was performed first using an End-it DNA End-Repair Kit (Epicenter, ER0720) according to manufacturer’s instructions, followed by DNA ligase-based adapter ligation (NEB, M2200S) and enrichment of adapter-modified DNA fragments by PCR. PCR fragments were gel-purified using a Qiagen Gel Purification Kit (Qiagen) and sequenced on an Illumina single-end flow cell. Sequencing of libraries was performed on Illumina HiSeq 2500v4 high-throughput sequencer. Fastq read files were trimmed using Cutadapt[DOI:10.14806/ej.17.1.200]. The trimmed reads were mapped to mm10 mouse genome using HISAT2 [PMID: 31375807]. Peak calling was conducted by HOMER program run in factor mode (FDR< 0.001). The intergenic peaks nearest to a TSS is annotated as the corresponding gene, and peaks within a gene were further categorized into exonic, intronic, 3’ or 5’ UTR regions using HOMER mm10 annotation. Venn diagrams were generated from DEG lists using IteractiVenn (interactivenn.net) (Heberle et al., 2015).

### Fluidigm qRT-PCR

Fluidigm qRT-PCR analysis was performed using previously published protocols (Hernandez et al., 2020; Pietras et al., 2016; Rabe et al., 2020). Pools of 100 cells were sorted directly into a 96-well PCR plate (AB2396, ThermoFisher) containing 5μL of 2x CellsDirect Reaction Mix (Invitrogen). Immediately following sort, the plate was sealed, centrifuged at 500 x g for 5 minutes, snap-frozen, and stored at −80°C. cDNA was generated from RNA by reverse transcription using Superscript III *Taq* polymerase (Invitrogen) and pre-amplified with a custom 96-target DeltaGene (Fluidigm) primer set for 18 cycles on a thermocycler (Eppendorf). Any excess primers from the pre-amplification reaction were removed by Exonuclease-I (New England Biolabs) incubation and samples were diluted in DNA suspension buffer (Teknova) prior to chip loading. A Fludigm 96.96 Dynamic Array IFC was loaded with cDNA, primers and SsoFast Sybr Green Master Mix (BioRad) and analyzed on a BioMark HD system (Fluidigm). Subsequently, data were analyzed using Fluidigm Gene Expression Software and all values were normalized to *Gusb*. The ΔΔCt approach was used to identify relative changes in gene expression. Hierarchical clustering and principal component analyses of normalized expression data were performed using ClustVis (biit.cs.ut.ee/clustvis) (Metsalu and Vilo, 2015). A complete list of qRT-PCR primers included in the custom primer sets as well as their sequences can be found in **Table S6**. Primer sets for *Hes1, Hoxa2, Hmga1, Ms4a3, Lcn2* and *Sod2* performed poorly due to low expression levels in HSC^LT^ and resulting data were not included in downstream analyses.

### Western blotting

2×10^4^ HSC were double-sorted into 1.5 ml tubes and centrifuged at 500 x g for 5 min in a swinging-bucket rotor. Supernatant was removed save for 100 μl. Cells were resuspended in this volume, transferred to 0.2 ml PCR tubes and centrifuged again at 500 x g for 5 min. Supernatant was carefully removed to leave 10 μl left in each tube. 10 μl PBS containing protease inhibitor cocktail (Roche) was added to each tube and cells were resuspended. 10 μl 4x Laemmli buffer (BioRad) was added to each tube, followed by incubation at 95°C for 10 min in a PCR cycler (Eppendorf). 30ul cell lysate was resolved on 4-20% SDS-PAGE precast gels (BioRad) at 100V. Protein was transferred to a PVDF membrane using a Trans-Blot semi-dry transfer system (BioRad). Following transfer, the membrane was blocked with 10 ml 1% bovine serum albumin (BSA) in TBS-T (TBS with 0.1% Tween 20) 1 hour at room temperature and subsequently incubated with anti-Myc or anti-Histone 3 antibody on a shaker overnight at 4°C. The membrane was then washed 5 times, 5 min for each wash in TBS-T at RT, incubated with secondary antibody at RT on a shaker for 1 hour, washed 5 times in TBS-T at room temperature and signals were detected using West Pico Plus ECL detection reagent (Thermo Scientific). Chemiluminescence was subsequently imaged on a BioRad gel imager.

### Statistical analysis

Statistical analyses were performed using Prism software (GraphPad). For RNA-seq, deSeq2 was used. Means and S.D. are reported except where noted. In figure legends, n refers to the number of biological replicates. The number of independent experiments from which the replicates derive is also reported. Except for RNA-seq and ChIP-seq analyses, statistical significance was determined by Mann-Whitney *u*-test (bivariate comparisons) or oneway ANOVA with Tukey’s test (multivariate comparisons).

## List of Supplemental Tables

**Table S1, related to Figure 1**. Differentially expressed genes in SLAM cells from mice treated ± IL-1β for 20 days.

**Table S2, related to Figure 1**. GO categories, Ingenuity Pathway Analysis Upstream Regulator categories and GSEA enrichment analysis of differentially expressed genes in SLAM cells from mice treated ± IL-1β for 20 days.

**Table S3, related to Figure 3**. Overlap between IL-1 downregulated genes and public datasets.

**Table S4, related to Figure 4**. PU.1 ChIP-seq peak locations, scores and PU.1/Myc motif scores.

**Table S5, related to Figure 4**. GO categories enriched in IL-1 DEGs containing PU.1 ChIP-seq peaks.

**Table S6, related to Methods**. Antibodies, antibody dilutions and Fluidigm qRT-PCR primers used in this study.

