## Supplemental Figures for "PU.1 enforces quiescence and limits hematopoietic stem cell expansion during inflammatory stress"

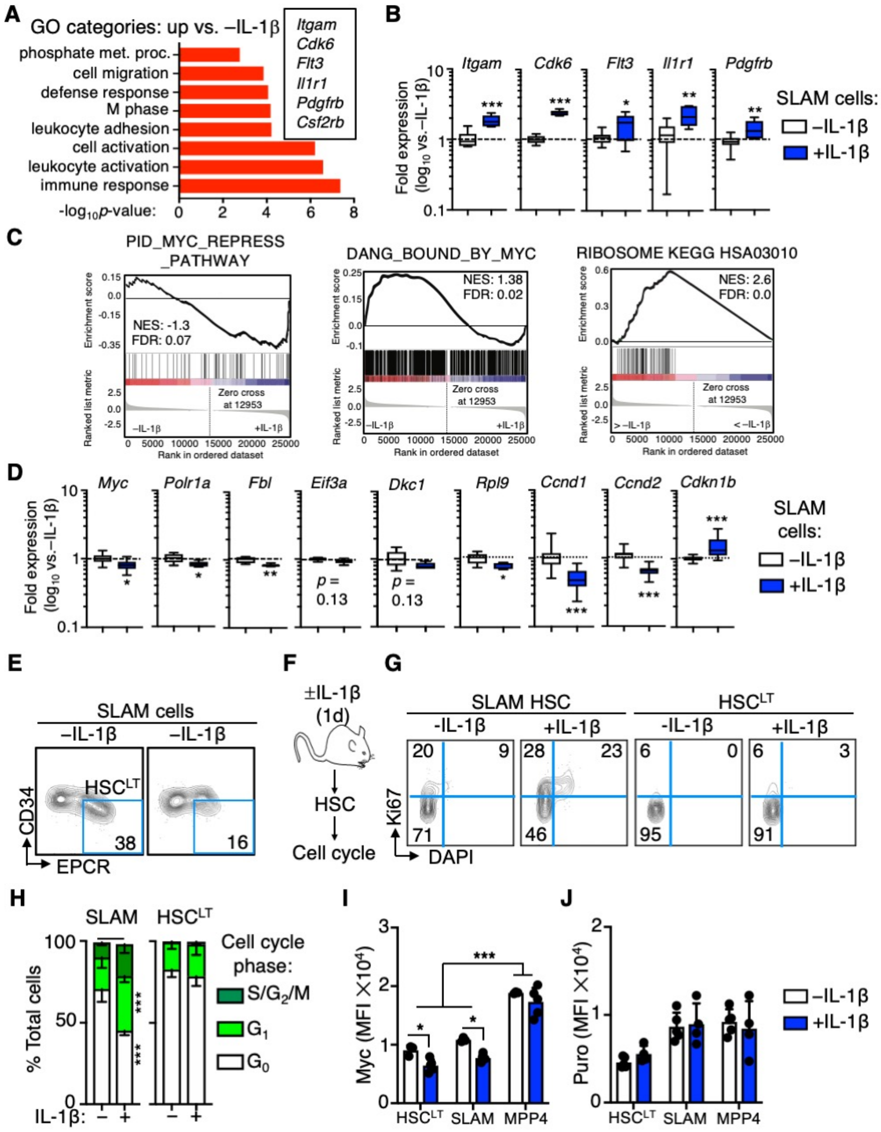


**Figure S1, related to Figure 1**. **A**) GO category enrichment of significantly upregulated DEGs in SLAM cells from mice treated 20d +IL-1β versus 20d -IL-1β, expressed as -$\log_{10}$ p-value. See Table S2 for complete list of GO categories. **B**) Quantification by Fluidigm qRT-PCR array of gene expression in SLAM cells from mice treated ± IL-1β for 20d (n = 8/group). Data are expressed as $\log_{10}$ fold expression versus -IL-1β. Box represents upper and lower quartiles with line representing median value. Whiskers represent minimum and maximum values. Data are representative of two independent experiments. **C**) Additional GSEA analysis of downregulated DEGs in SLAM cells from mice treated 20d +IL-1β versus 20d -IL-1β shown in Figure 1E. **D**) Quantification by Fluidigm qRT-PCR array of cell cycle and protein synthesis gene expression in SLAM cells from mice treated ± IL-1β for 20d (n = 8/group). Data are expressed as $\log_{10}$ fold expression versus -IL-1β. Box represents upper and lower quartiles with line representing median value. Whiskers represent minimum and maximum values. Data are representative of two independent experiments. **E**) Representative FACS plots showing frequencies of phenotypic $HSC^{LT}$ fraction within the SLAM gate from mice treated 20d +IL-1β versus 20d -IL-1β. **F**) Experimental design, **G**) Representative FACS plots and **H**) quantification of cell cycle phase distribution in SLAM cells and $HSC^{LT}$ (n = 5/group). **I**) Geometric mean fluorescence intensity

(MFI) of Myc from one representative experiment (n = 5/grp). Individual values are shown with bars representing mean values. **J**) Geometric mean fluorescence intensity (MFI) of Puro from one representative experiment (n = 5/grp). Individual values are shown with bars representing mean values. * $p < 0.05$, ** $p < 0.01$, ***$p < 0.001$ by Mann-Whitney *u*-test or one-way ANOVA with Tukey's post-test. Error bars represent S.D.

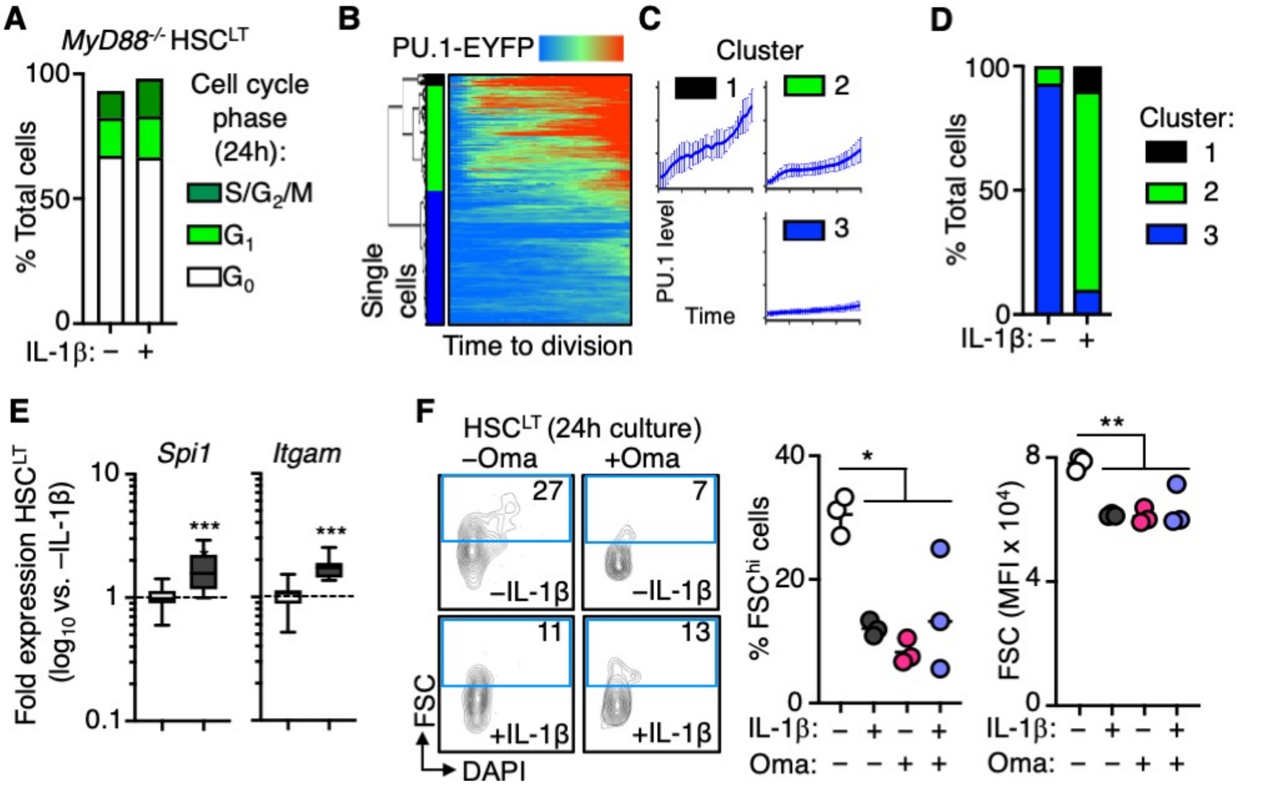


**Figure S2, related to Figure 2**. **A**) Quantification of cell cycle phase distribution in HSC$^{LT}$ treated *in vitro* ± IL-1β for 24h based on Ki67 and DAPI (n = 2/group). Data are from one experiment. **B**) Hierarchical clustering and heatmap showing PU.1-EYFP expression over time prior to first division in individual HSC$^{LT}$ treated *in vitro* ± IL-1β. Data represent one experiment. **C**) PU.1 expression over time in HSC$^{LT}$ based on experiment in B). **D**) Frequency of HSC$^{LT}$ in each cluster based on experiment in B). **E**) Quantification by Fluidigm qRT-PCR array of cell cycle and protein synthesis gene expression in HSC$_{LT}$ from mice treated ± IL-1β for 12h (n = 16/group). Data are expressed as log$_{10}$ fold expression versus -IL-1β. Box represents upper and lower quartiles with line representing median value. Whiskers represent minimum and maximum values. Data are compiled from two independent experiments. **F**) Representative FACS plots (left), quantification of FSC$^{hi}$ HSC based on FSC/DAPI content (center) and quantification of FSC geometric mean fluorescence intensity (MFI; right) in HSC$^{LT}$ treated *in vitro* ± IL-1β or ± Omacetaxine (Oma) for 24h. Individual values are shown with lines representing mean values. * $p < 0.05$, ** $p < 0.01$, *** $p < 0.001$ by Mann-Whitney *u*-test or one-way ANOVA with Tukey's test. Error bars represent S.D.

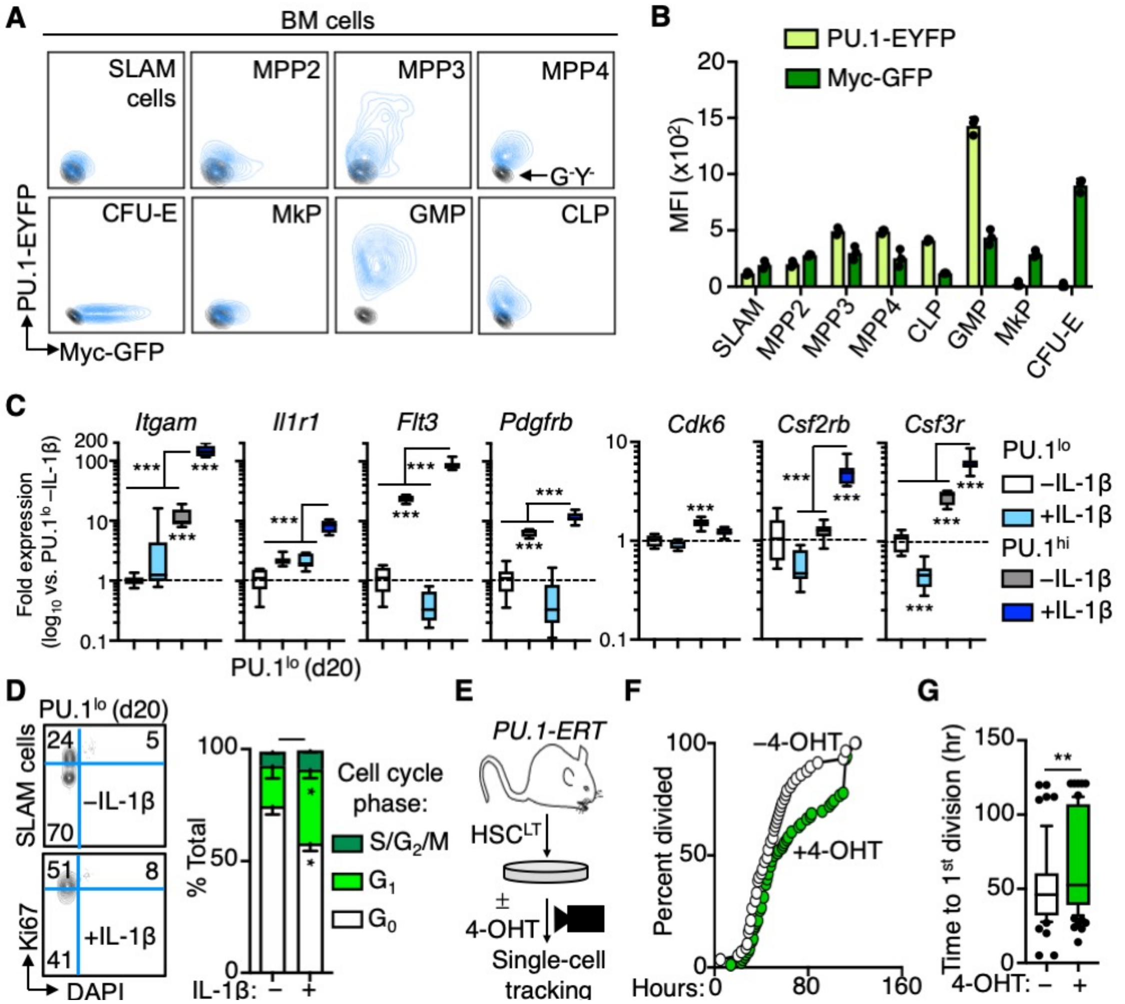


**Figure S3, related to Figure 3. A**) Representative FACS plots showing expression of Myc-GFP and PU.1-EYFP in the indicated hematopoietic populations (blue contours). GFP-/EYFP- control (G-/Y-) is shown in each plot (black contours). **B**) Quantification of geometric mean fluorescence intensity (MFI) of Myc-GFP and PU.1-EYFP in hematopoietic populations indicated in A) (n = 3/group). Individual values are shown with bars representing mean values. **C**) Quantification by Fluidigm qRT-PCR array of IL-1-upregulated gene expression in Pu.$1^{lo}$ and PU.$1^{hi}$ SLAM fractions from *PU.1-EYFP* mice treated ± IL-1β for 20d (n = 8/group). Data are expressed as $log_{10}$ fold expression versus -IL-1β. Box represents upper and lower quartiles with line representing median value. Whiskers represent minimum and maximum values. Data are representative of two independent experiments. Representative FACS plots (left) and quantification (right) of cell cycle distribution in PU.$1^{lo}$ SLAM cells from mice treated ± IL-1β for 20d (n = 3/group) using Ki67 and DAPI. Data are compiled from two independent experiments. **E**) Experimental design for single cell tracking studies of *PU-ER* $HSC^{LT}$ cultured ± 4-OHT. **F**) Graph showing kinetics of first cell division in $HSC^{LT}$ cultured ± IL-1β (n = 194 -IL-1β, 139 IL-1β). Data are compiled from three independent experiments. **G**) Cumulative time to first cell division (n = 57 -4-OHT, 86 +4-OHT). Data are from one experiment. Box shows upper and lower quartiles with line showing median value; whiskers upper and lower $10^{th}$ percentile and individual dots represent outliers. ***p< 0.001 by one-way ANOVA with Tukey's test. Error bars represent S.D.

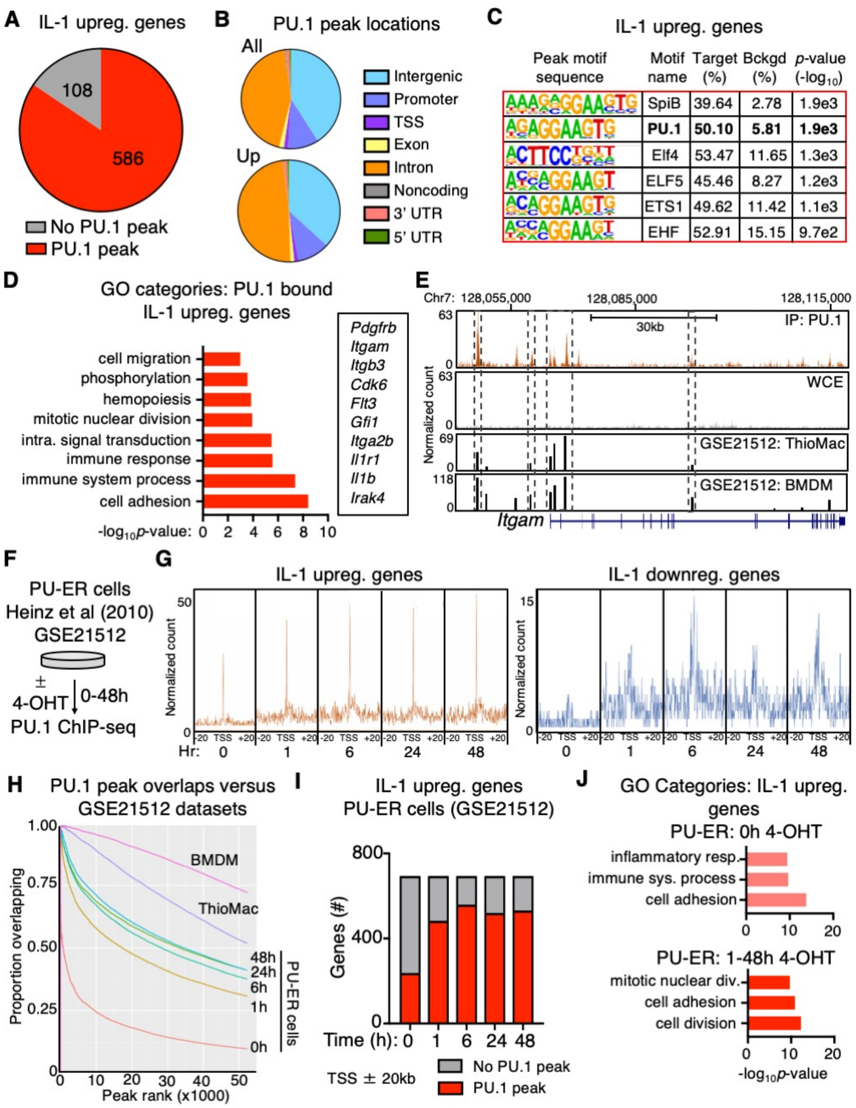


**Figure S4, related to Figure 4**. **A**) Pie chart comparing IL-1 upregulated genes identified in SLAM cells by RNA-seq analysis in Figure 1D and presence of PU.1 peaks at or near these genes (TSS ± 20 kb) in ChIP-seq data. See also Table S7. **B**) Pie chart showing proportion of PU.1 peak locations in all genes versus IL-1 upregulated genes. **C**) Transcription factor binding site motif enrichment at PU.1 ChIP-seq peak sites located at TSS ± 20 kb in IL-1-downregulated genes. **D**) GO category enrichment of IL-1 upregulated DEGs containing PU.1 peaks. Representative genes in the indicated categories are shown to the right. Data are expressed as -$\log_{10}$ *p*-value. See also Table S9. **E**) UCSC genome browser rendering of PU.1 peak location in *Itgam* gene body. Tracks show PU.1 ChIP-Seq, whole cell extract (WCE) control, and peak locations and intensities in thyoglycollate-elicited primary mouse macrophage (ThioMac) and bone marrow-derived macrophage (BMDM) PU.1 ChIP-Seq datasets from GSE21512. **F**) Experimental design used for generation of PU-ER dataset in GSE21512. **G**) PU.1 peak locations and intensities in IL-1 up and downregulated genes in PU-ER cells (TSS ± 20 kb). **H**) Comparison of PU.1 peak overlaps between datasets **I**) Comparison of IL-1 upregulated genes in SLAM HSC with genes containing PU.1 peaks within TSS ± 20 kb in PU-ER cells ± 4-OHT. Based on PU.1 ChIP-seq datasets in GSE21512. **J)** GO category enrichment of IL-1 downregulated DEGs containing PU.1 peaks in PU-ER cell ChIP-seq dataset at 0h +4-OHT versus combined 1-48h +4-OHT. Top three GO categories are shown. Data are expressed as -$\log_{10}$ *p*-value.

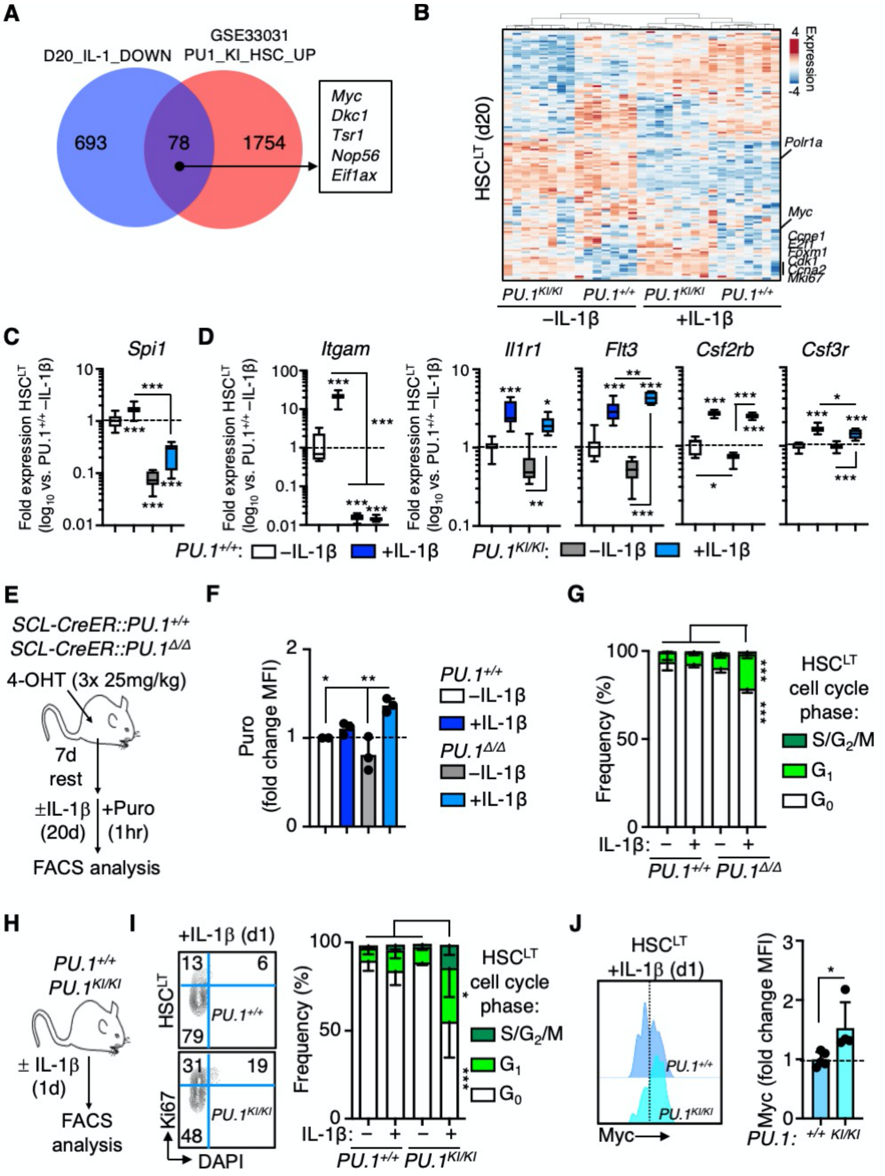


**Figure S5, related to Figure 5**. **A**) Venn diagram comparing genes downregulated by IL-1 in SLAM cells and genes upregulated in $PU.1^{KI/KI}$ SLAM cells. **B**) Heatmap and hierarchical clustering analysis (Pearson Correlation) of Fluidigm qRT-PCR analyses in Figure 5B. **C-D**) Quantification by Fluidigm qRT-PCR array of cell cycle and protein synthesis gene expression in HSC$^{LT}$ in Figure 5B. Data are expressed as log$_{10}$ fold expression versus -IL-1β. Box represents upper and lower quartiles with line representing median value. Whiskers represent minimum and maximum values. **E**) Study design for Cre induction with 4-OHT and analysis of HSC$^{LT}$ from *SCL-CreERT* $PU.1^{+/+}$ and $PU.1^{Δ/Δ}$ mice treated ± IL-1β for 20d (n = 2-3/group). **F**) Intracellular flow cytometry analysis of puromycin (Puro) incorporation in HSC$^{LT}$ from mice in Figure S5E. Puro was injected i.p. 1 hour before BM harvest. Data are expressed as fold change of mean fluorescence intensity (MFI) versus -IL-1β. Individual values are shown with bars representing mean values. Data are from one experiment. **G**) Quantification of cell cycle phase distribution in HSC$^{LT}$ from $PU.1^{+/+}$ and $PU.1^{Δ/Δ}$ mice in Figure S5E based on Ki67 and DAPI. Data are from one experiment **H**) Experimental design for analysis of HSC$^{LT}$ from $PU.1^{+/+}$ and $PU.1^{KI/KI}$ mice treated ± IL-1β for 1d (n = 3-5/group). **I**) Representative FACS plots (left) and quantification (right) of cell cycle phase distribution in $HSC^{LT}$ from mice in Figure S5H based on Ki67 and DAPI. Data are from one experiment. **J**) Myc levels from *PU.1*$^{+/+}$ and *PU.1*$^{KI/KI}$ mice treated +IL-1β for 1d (n = 4/group). Individual values are shown with bars representing mean values. Data are from one experiment. * $p < 0.05$, ** $p < 0.01$, ***$p < 0.001$ by Mann-Whitney *u*-test or one-way ANOVA with Tukey's test. Error bars represent S.D.

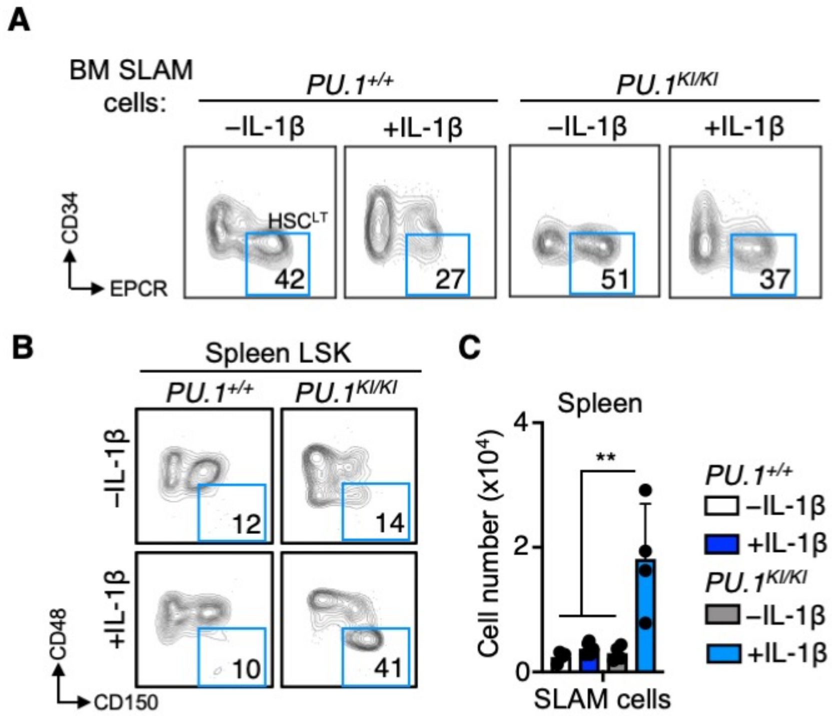


**Figure S6, related to Figure 5. A**) Representative FACS plots showing gating strategy for identification of $HSC^{LT}$ from BM of *PU.1*$^{+/+}$ and *PU.1*$^{KI/KI}$ mice treated ± IL-1β for 20d **B**) Representative FACS plots showing gating strategy for identification of SLAM cells from spleens of *PU.1*$^{+/+}$ and *PU.1*$^{KI/KI}$ mice treated ± IL-1β for 20d. **C**) Quantification of SLAM cells from spleens of *PU.1*$^{+/+}$ and *PU.1*$^{KI/KI}$ mice treated ± IL-1β for 20d (n = 4/group). Data are from one experiment. * $p < 0.05$, ** $p < 0.01$, ***$p < 0.001$ by one-way ANOVA with Tukey's test. Error bars represent S.D.
